# Thymocyte regulatory variant alters transcription factor binding and protects from type 1 diabetes in infants

**DOI:** 10.1101/2021.09.17.460789

**Authors:** Niina Sandholm, Arcadio Rubio García, Marcin L Pekalski, Jamie RJ Inshaw, Antony J Cutler, John A Todd

## Abstract

We recently mapped a genetic susceptibility locus on chromosome 6q22.33 for type 1 diabetes diagnosed below age of 7 years near the gene Protein tyrosine phosphatase receptor type K (*PTPRK)* and the thymocyte selection associated gene (*THEMIS)*. As the thymus plays a central role in shaping the T cell repertoire, we aimed to identify the most likely causal genetic factors behind the association using thymocyte genomic data. In four thymocyte populations we identified 253 DNA sequence motifs underlying histone modifications. The G insertion allele of rs138300818, associated with protection from diabetes, created thymocyte motifs for multiple histone modifications and thymocyte types. The insertion also disrupted a predicted RFX5/7 transcription factor binding site. RFX7 is abundantly expressed in thymus. Chromatin state and RNA sequencing data suggested strong transcription overlapping rs138300818 in fetal thymus, while eQTL and chromatin conformation data indicated that the rs138300818 insertion is associated with lower *THEMIS* expression. Taken together, our results support a role for thymic *THEMIS* gene expression and the rs138300818 variant in promoting the development and diagnosis of type 1 diabetes at an earlier age.

---

Over 1 million children and adolescents have type 1 diabetes, a disease caused by an autoimmune reaction against the insulin-producing beta cells in the pancreatic islets. The HLA class II and I genes, which present pathogen and self-antigens to T cells, play an important role in development of type 1 diabetes and other autoimmune diseases; in addition, over 100 other genetic loci have been associated with type 1 diabetes (1–4). Including the most susceptible HLA class II genotype, DRB1*03-DQB1*02/DRB1*0401-DQB1*0302, several of the non-HLA gene variants lower the age-of-diagnosis (AAD) of type 1 diabetes (5).

The HLA proteins bind antigen fragments, peptides, which are then recognized by highly variable T cell antigen receptors (TCR), found on the T cell surface. In the thymus, the T cell precursors, called thymocytes, are first matured from CD4^−^CD8^−^ double negative cells to CD4^+^CD8^+^ double positive cells. In positive selection, thymocytes are limited to those with sufficient TCR binding to the HLA proteins; the cells with affinity to HLA class II proteins (found on antigen presenting cells) differentiate towards CD4^+^ T-helper cells; and the ones with affinity to HLA class I (expressed by most cells) differentiate towards CD8^+^ T-killer cells. Negative selection excludes the T cells targeting the self, thus protecting from autoimmunity (6). Therefore, thymus has been suggested to play a role in development of type 1 diabetes and other autoimmune diseases (7,8). Support for this hypothesis came from a genome-wide association study (GWAS) on type 1 diabetes where the susceptibility loci were enriched on active enhancer sites in thymus, in addition to T, B and NK cells and CD34^+^ stem cells (9). Furthermore, a recent single-cell RNA sequencing (scRNA-seq) study on developing mouse thymus identified a modest enrichment of GWAS-signals from type 1 diabetes and other autoimmune diseases especially in the thymus blood cell populations (10).

The thymus plays an important role in training the immune system in early childhood when the body is adapting to the changing environment; the thymus itself is proportionally largest at birth, and nearly disappears in adults. Therefore, the role of thymus in autoimmunity is likely most pronounced for the childhood autoimmune diseases, including a key role the infant gut microbiome, where we have discovered a cross-reactive mimotope of the primary autoantigen in type 1 diabetes, insulin, encoded in the commensal bacterial enzyme, transketolase (11). Recently it has been shown in mice that intestinal dendritic cells can engulf gut bacteria and traffic to the thymus where the bacterial antigens are presented to the T cell immune system (12). Hence, expression of insulin in the human thymus is protective for type 1 diabetes and now expression of bacterial antigens early in life in the thymus adds to the body's attempts to establish and maintain immune tolerance to insulin.

A recent genetic analysis of AAD of type 1 diabetes identified a novel locus on chromosome 6q22.33 with variants associated with earlier diagnosis of type 1 diabetes (rs72975913 p=2.94×10^−10^) (13); the locus was associated with diabetes particularly among those diagnosed in the first 7 years (5). The association is located between the thymocyte selection associated (*THEMIS*) and Protein tyrosine phosphatase, receptor type K (*PTPRK*) genes. *THEMIS* plays a regulatory role in both positive and negative T cell selection during late thymocyte development and is necessary for proper lineage commitment and maturation of T cells. While *PTPRK* is widely expressed and regulates a variety of cellular processes including cell growth and differentiation, *PTPRK* has also been shown to regulate CD4^+^ positive T cell development (14).

Variants associated with common diseases are enriched in transcription factor (TF) binding sites and active chromatin regions, suggesting a regulatory role for the disease outcome (15). In this study, we hypothesized that the association at the recently identified 6q22.33 locus is mediated through regulatory processes, and used extensive genomic data from thymus and other tissues to pinpoint the likely causal variants and the functional mechanisms underlying the association.

## RESEARCH DESIGN AND METHODS

### Overall study design

We sought DNA motifs located in active chromosome regions of thymocyte cells, defined as histone methylation and overlapping RNA sequencing (RNA-seq) expression data. Resulting thymocyte motifs were integrated with the 58 credible set SNPs on the 6q22.33 locus, and 1,165 other SNPs for type 1 diabetes, to identify SNPs that create DNA sequences matching the thymocyte motifs (**Figure 1**).

**Figure 1:**
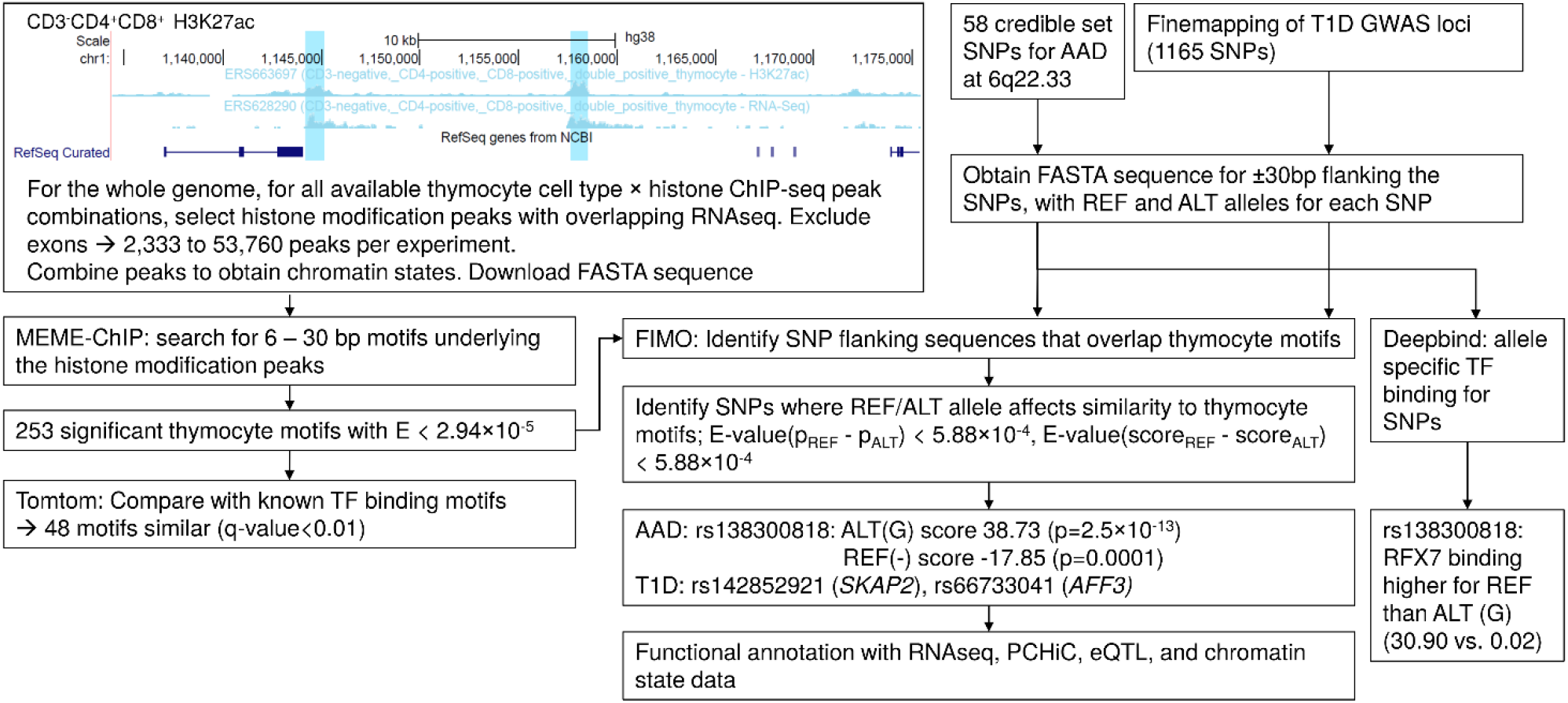
Schematic illustration of the study design.

### Study material

Genome-wide chromatin immunoprecipitation sequencing (ChIP-seq) data on histone modification marks were available from the Blueprint epigenomics project (16) (accessed through ftp://ftp.ebi.ac.uk/pub/databases/blueprint/data/homo_sapiens/GRCh38/) for four individuals. Available peaks included H3K4 trimethylation (H3K4me3), H3K4 monomethylation (H3K4me1), H3K27 acetylation (H3K27ac), H3K36 trimethylation (H3K36me3), and H3K27 trimethylation (H3K27me3). Samples were dissected to CD3^−^CD4^+^CD8^+^ thymocytes, CD3^+^CD4^+^CD8^+^ thymocytes, CD4^+^αβ thymocytes, and CD8^+^αβ thymocytes. Total RNA-seq data was available and downloaded for these cell types for one individual.

Human reference genome DNA sequence primary assembly, and exon annotations were downloaded from Ensembl (ftp.ensembl.org/pub/release-92/) for human genome build GRCh38, release 92. DNA positions were updated from human genome build GRCh37 to GRCh38 using UCSC liftover tool (https://genome.ucsc.edu/cgi-bin/hgLiftOver) when necessary.

SNPs associated with type 1 diabetes were obtained from Burren et al. (17), where credible set SNPs were defined as those with group posterior probability ≥0.9 from GUESSFM statistical fine mapping of ImmunoChip GWAS data, further filtered to a total of 1,165 variants with p-value < 1×10^−5^ for association with type 1 diabetes.

RNA expression for the genes of interest was explored with the Roadmap epigenome browser (https://epigenomegateway.wustl.edu/browser) for available blood cells (primary mononuclear cells, CD4^+^ Memory Primary cells, CD4^+^ naive primary cells, primary CD8^+^ naïve T cells) and thymus. Furthermore, we queried RNA expression data from Human Protein Atlas (HPA) version 18, for HPA, Genotype-Tissue Expression (GTEx), and Functional Annotation of Mammalian Genomes 5 (FANTOM5) transcriptome data sets.

3D chromatin conformation capture data based on Promoter capture Hi-C (PCHiC) were obtained using CHiCP (18) for 16 primary blood cell types and for thymus (19), for CD34^+^ stem cells and for the human Epstein-Barr virus (EBV)-transformed lymphoblastoid cell line GM12878 (20), and for pancreatic islets (21). Interactions with CHiCAGO score > 5 were considered high-confidence (19). eQTL associations were queried in the GTEX (any tissue, gtexportal.org/) and eQTLgen whole blood cis-eQTL data bases (22).

### Histone modification ChIP-seq peak selection and quality control

bedtools v 2.27.1 software package (23) and intersect tool were used for processing the histone modification peaks. Histone modification peaks were limited to those with peak fold change ≥3 and q-value < 0.0001. To enrich the search for active DNA sites, we limited the ChIP-seq peaks to those with total RNA-seq overlapping the peak, defined as mean RNA-seq depth ≥0.001 over the peak length, using UCSC bigWigSummary and bigWigAverageOverBed utilities (http://hgdownload.soe.ucsc.edu/admin/exe/). Finally, overlapping peaks of the same ChIP-seq mark and cell type, but from different biological repeats (individuals), were pooled together **(Supplementary Table S1)**.

Based on these pooled histone modification peaks, we defined cell type specific chromatin states as intersect of the contributing histone marks using bedtools intersect tool, mimicking a previous definition by Ernst and Kellis (24) for state7 (H3K4me1 and H3K36me3), state9 (H3K4me1 and H3K27Ac), state10 (H3K4me3, H3K27Ac, and H3K4me1), state11 (H3K4me3 and H3K4me1), and state12 (H3K4me3 and H3K27Ac). States 7, 9, 10, and 11 were available only for the CD3^+^CD4^+^CD8^+^ thymocytes, whereas state 12 was additionally available for the CD3^−^CD4^+^CD8^+^ and CD4^+^αß thymocytes **(Supplementary Table S2)**. Other states were either based on single histone mark and thus already covered, or contained other histone marks not available for the thymocytes. Peak calls shorter than six base pairs (bp) were excluded. A flanking region of 20 bp was then added to allow TF binding motifs on the peak borders, and exonic regions were excluded.

### DNA Motif search

Fasta sequences were extracted for each peak with bedtools getfasta tool. Motif discovery was performed using MEME Suite (25) running MEME-ChIP, searching for 6 to 30bp motifs enriched relative to a random first order model (i.e. adjusting for nucleotide and dimer biases, e.g. CG content), targeting 20 motifs. As by default, 600 sequences were randomly selected for initial motif discovery. Significant motifs were defined as E-value < 0.01/17 cell type and histone mark combinations/20 motifs = 2.94×10^−5^. DREME search for short motifs was performed until E-value<0.05. Resulting motifs were compared to the *in vivo* and *in silico* vertebrate DNA motifs (including JASPAR CORE vertebrates, UniProbe Mouse, and Jolma2013 Human and Mouse data bases) with tomtom tool (25); significant similarity was defined as q-value < 0.01.

### Motif and sequence overlap

Fasta sequences were extracted for ±30 bp around the 58 credible set SNPs for both the reference and alternative allele from ensembl (build GRCh38) using R jsonlite package v.1.5 (26). Sequence overlap with the discovered thymocyte motifs was studied with FIMO tool from the MEME suite (25). Significant motif overlap was considered as P_REF_ or P_ALT_ < 0.01/253 significant thymocyte motifs (from MEME)/58 SNPs = 6.81×10^−7^.

The thymocyte motif overlap was calculated similarly for SNPs in the credible sets associated with type 1 diabetes (17). Significant motif overlap was considered as P_REF_ or P_ALT_ < 0.01/253 significant thymocyte motifs/1,165 SNPs = 3.39×10^−8^.

We calculated an E-value (pREF - pALT) for the difference in *p*-value between the alleles by calculating a z-score as (mean difference in p-values – difference in p-values)/(standard deviation of differences in p-values), and converting the z-score into an E-value (p-value) assuming two-tailed normal distribution; this score was conservative, as only those SNP – motif pairs were considered where at least one of the SNP alleles had nominally significant (p<0.05) similarity with a motif sequence. Distributions were calculated separately for each cell-type – histone modification combinations (N=17), and E-value(p_REF_ - p_ALT_) < 0.01/17 = 5.88×10^−4^ was considered significant.

### Allele-specific transcription factor binding sites

We employed DeepBind v0.11 (27) to assess allele-specific TF binding for the 58 SNPs in the credible set for AAD. DeepBind was run using default parameters, and all non-deprecated TFs delivered with the distribution (N=927).

### Data and Resource Availability

All utilised data were downloaded from publicly available data bases. All utilised software is freely available for research purposes.

## RESULTS

Statistical fine mapping of the 6q22.33 locus previously suggested 58 SNPs in the credible set of likely causal variants across three LD blocs; the posterior probability of the underlying causal variant being among the three groups was estimated 0.96, 0.50, and 0.42, respectively (13). PCHiC data, which identifies DNA fragments in contact with gene promoter regions, indicated that many of the 58 SNPs were located in fragments that interact with *PTPRK* (in foetal thymus, naïve CD4^+^ T cells) and *THEMIS* (in Naïve CD8^+^ T cells) promoter regions (ChiCAGO score ≥5 (19); **Supplementary Table 3**). The SNPs interacted with no other genes in thymus or in the 16 primary hematopoietic cells, and as a negative control, they had no PCHiC interactions in the pancreatic islets (21). The strongest gene expression of *THEMIS* is found in thymus in the FANTOM5 database (28), and it is also expressed in T cells; also *PTPRK* and three other genes within 1 Mbp from the lead SNP rs72975913 had detectable expression in thymus (**Supplementary Table S4**).

### Active DNA sequence motifs in thymocytes

With the hypothesis that the causal mechanism of the association signal is through gene regulation in thymocytes, we set to search for regulatory DNA motifs in thymocytes, affected by the credible set of SNPs. Genome-wide histone modification marks, including H3K4me3, H3K4me1, H3K27ac, H3K36me3, and H3K27me3, were available for thymocytes that were dissected to four cell types (CD3^−^CD4^+^CD8^+^, CD3^+^CD4^+^CD8^+^, CD4^+^αβ, and CD8^+^αβ). Number of peaks ranged from 7,088 to 55,622 per experiment **(Supplementary Table S1)**. Data from different individuals was then pooled over the four cell types to mimic previously reported chromatin states consisting of combination of histone modification marks (29) (**Supplementary Table S2**).

Search for DNA sequence motifs reoccurring within these histone modification and chromatin state peaks, indicative of active chromatin in corresponding thymocyte cells, resulted in 253 motifs (E-value < 2.94×10^−5^) ranging from one to 20 motifs identified for different cell type and peak specific data sets (**Table 1**). Among these, 48 motifs were similar (*q*-value<0.01) to known binding motifs for SP1, SP2, SP3, SP4, KLF5, KLF16, ZNF263, mouse Zfp281 (human ortholog ZNF281), mouse Zfp740 (human ortholog ZNF740), Zfx, and RREB1 **(Supplementary Table S5, Supplementary Figure S1)**. While most of these TFs are expressed in nearly all tissues, it is of note that SP3 has the highest expression in thymus across the FANTOM5 tissues (159.9 transcripts per million [tpm]), along with high thymus expression of ZFP281 (43.2 tpm) as well.

**Table 1:** Number of histone modification peaks in genome, and identified motifs per cell type.

| Histone mod | Cell type | N samples | N peaks | N motifs |
| --- | --- | --- | --- | --- |
| H3K4me3 | CD3<sup>+</sup>CD4<sup>+</sup>CD8<sup>+</sup> | 3 | 48,295 | 17 |
|  | CD3<sup>-</sup>CD4<sup>+</sup>CD8<sup>+</sup> | 2 | 33,132 | 19 |
| | CD4<sup>+</sup>$\alpha\beta$ | 1 | 37,452 | 20 |
| | CD8<sup>+</sup>$\alpha\beta$ | 1 | 36,825 | 18 |
| H3K4me1 | CD3<sup>+</sup>CD4<sup>+</sup>CD8<sup>+</sup> | 1 | 36,308 | 2 |
| H3K27ac | CD3<sup>+</sup>CD4<sup>+</sup>CD8<sup>+</sup> | 2 | 33,131 | 18 |
|  | CD3<sup>-</sup>CD4<sup>+</sup>CD8<sup>+</sup> | 1 | 12,334 | 17 |
| | CD4<sup>+</sup>$\alpha\beta$ | 1 | 6,011 | 18 |
| H3K36me3 | CD3<sup>+</sup>CD4<sup>+</sup>CD8<sup>+</sup> | 2 | 53,760 | 1 |
| H3K27me3 | CD3<sup>-</sup>CD4<sup>+</sup>CD8<sup>+</sup> | 1 | 2,333 | 20 |

N samples: number of samples available for the cell type and histone modification. N peaks: number of peaks after pooling all peaks from the same cell type, and excluding exons. N motifs: Number of significant DNA motifs (E-value < $2.94 \times 10^{-5}$ ) underlying each cell type specific histone modification peak, identified with MEME software.

### Credible set SNPs affecting thymocyte motifs

Many of the obtained sequence motifs for active thymocyte regions overlapped the 58 AAD SNPs. For five out of the 58 SNPs, the flanking DNA sequences were similar to thymocyte motifs (*p* < 5.07×10^−7^ for REF or ALT) and had significantly different *p*-value for the REF and ALT alleles (E-value(*p*diff) < 5.88×10^−4^; **Supplementary Table S6**). The most marked allelic difference was observed for rs138300818 insertion variant for H3K27ac in the CD4^+^αß cells, with the score for motif-sequence similarity of 38.73 (*p* = 2.5×10^−13^) for the alternative G-insertion allele, and −17.85 (p=0.0001) for the reference allele (**Figure 2 A–C**). Of note, this was also the only variant with a significant allelic difference in score (E-value(Scorediff) < 5.88×10^−4^). A similar motif affected by the same rs138300818 substitution was identified in altogether 16 cell type and histone modification combinations, including all the four studied thymocyte cell types, and H3K27ac, H3K4me3, H3K27me3, and states 7 (genic enhancers), 9 (active enhancers), 10 (distal active promoters) and 12 (active transcription start site (TSS)).

**Figure 2:**
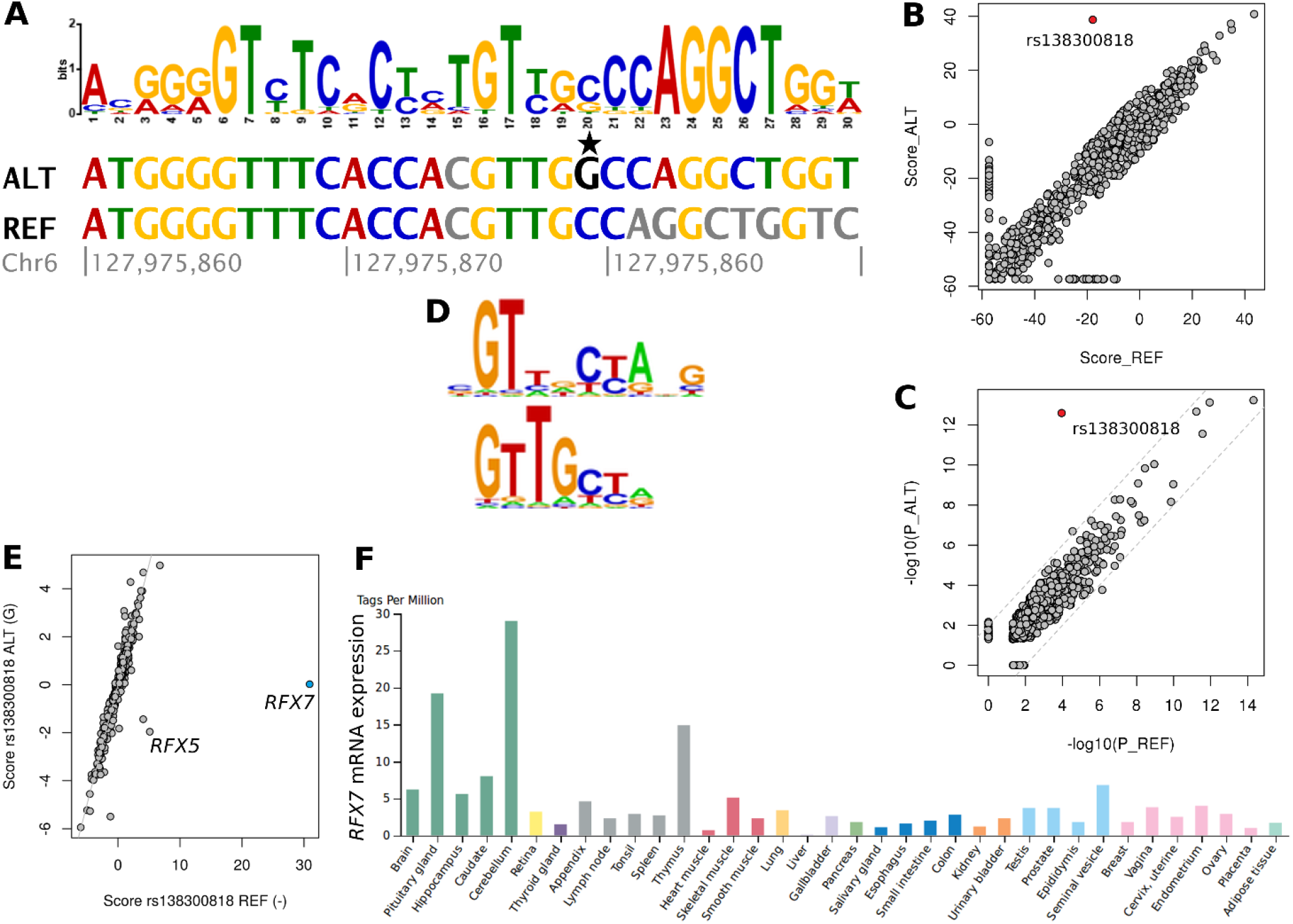
rs138300818 G insertion allele significantly increases sequence similarity with a thymocyte H3K27ac motif in CD4^+^αß cells. **A:** Overlap between a thymocyte H3K27ac motif and rs138300818 alternative G allele (highlighted with star) and reference null allele flanking sequence. Bases conflicting with the motif are indicated with gray color. **B and C:** Sequence similarity score and −log10(p-value) distributions (C) for all credible set SNP – motif pairs where at least one of the SNP alleles (REF or ALT) forms a motif overlapping sequence (p<0.05). rs138300818 is indicated as a red dot. **D:** rs138300818 REF allele, but not ALT insertion allele, forms a RFX7 transcription factor binding site. **E:** score distribution for rs138300818 REF and ALT alleles against all available transcription factor binding sites from DeepBind data base. **F:** RFX7 gene expression is highest in brain (cerebellum, pituitary gland), and in thymus in FANTOM5 data base.

The tomtom sequence similarity analysis did not find any known vertebrate TF-binding motifs similar to the thymocyte activity motifs overlapping rs138300818 (E > 0.05). However, a deep learning-based DeepBind method (27) identified RFX7 binding to the rs138300818 flanking sequence; the score for RFX7 binding was markedly higher for the REF null allele than for the ALT (G) allele (30.90 vs. 0.02, respectively; score mean 0.07, sd 1.39 for all transcripts; **Figure 2 D–F**). Of note, this represented the largest allelic difference in predicted TF binding across all credible set SNPs for AAD and available TFs (**Supplementary Figure S2**). Binding affinity was predicted higher for the reference allele also for two other RFX TF family members, RFX5 and RFX3 (**Supplementary Table S7**).

### Functional annotation for rs138300818 with PCHiC, eQTL, and chromatin state data

The variant rs138300818 has a minor allele frequency (MAF) of 0.155 (1000 Genomes [1000G] European population) for the G allele insertion, and it belongs to the highest 96% posterior probability group for association with AAD on the 6q22.33 locus (13). The variant is an intronic variant within the *PTPRK* gene. Roadmap fetal thymus ChromHMM model – based on histone modification data - predicted strong transcription overlapping rs138300818 (**Figure 3**). Indeed, non-coding RNA expression was detected overlapping rs138300818 in all four studied thymocyte cell types (**Figure 3**).

**Figure 3:**
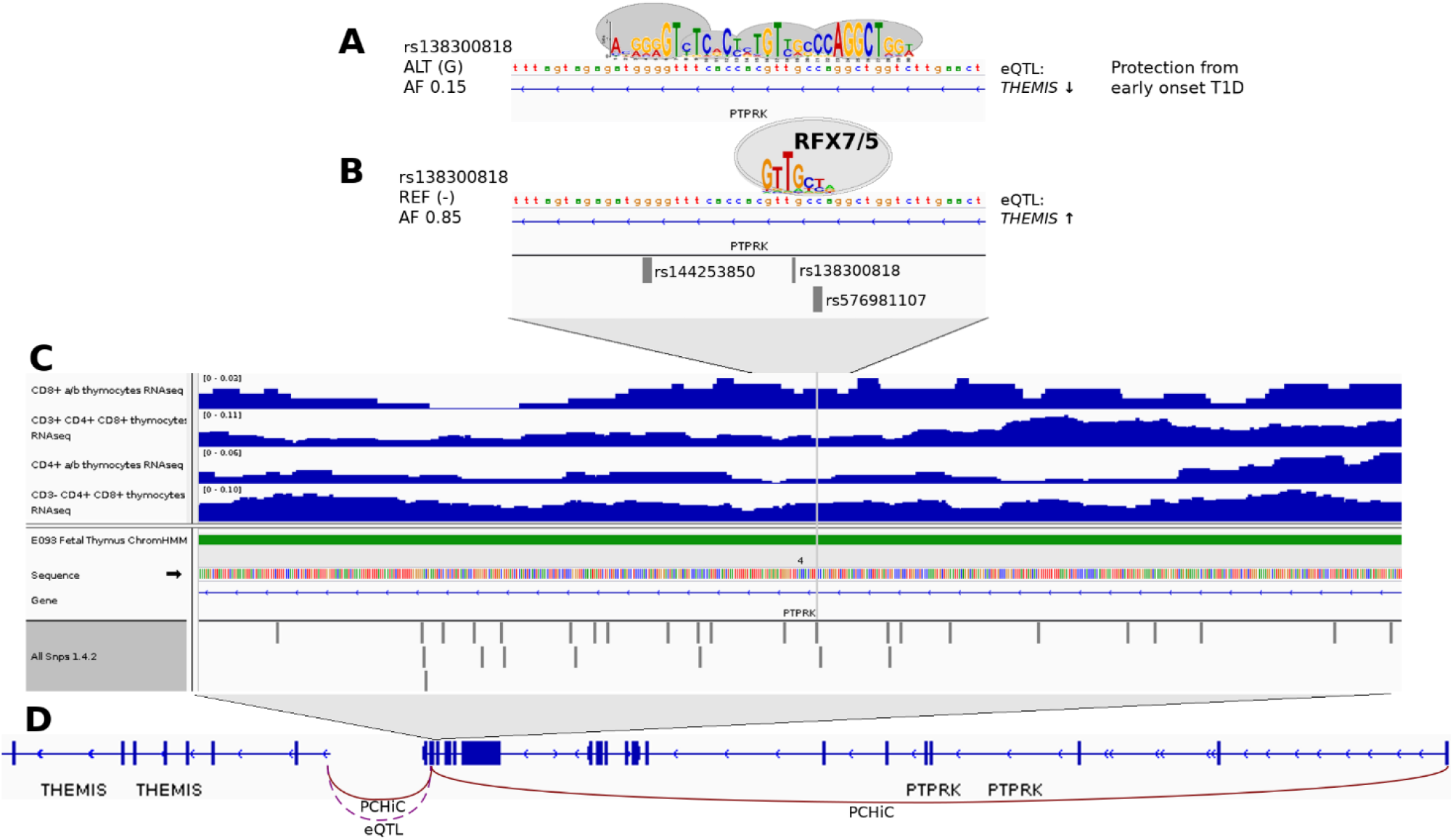
rs138300818 G insertion allele – protecting from early onset T1D – creates a thymocyte motif. **(A).** At the same time, this motif disrupts the RFX5/7 binding motif which is present with the rs138300818 reference (null) allele (**B**). rs138300818 is located intronic in *PTPRK*. Non-coding RNA transcription was detected overlapping rs138300818 (grey vertical line) in all four studied thymocyte cell types (dark blue peaks). Roadmap chromatin state data for fetal thymus (PrimaryHMM) indicated strong transcription (state 4, green bar) overlapping the SNP flanking region (**C**). PCHiC data in fetal thymocytes and in naïve CD8 cells suggests that rs138300818 interacts with both *PTPRK* and *THEMIS* transcription start sites. The genetic association signal for early onset T1D is colocalized with eQTL signal for *THEMIS* in human whole blood; rs138300818 reference allele – which forms the RFX7/5 binding motif – is associated with higher *THEMIS* expression (**D**).

PCHiC data for rs138300818 indicated interaction with *PTPRK* (CHiCAGO scores 13.79 and 1.81, in thymus and naïve CD8 cells, respectively), and with *THEMIS* (CHiCAGO scores 2.25 and 5.65, in thymus and naïve CD8 cells, respectively). Insertion/deletion variants (indels), e.g., rs138300818, are not included in the eQTLgen data base comprising of 31,684 whole blood samples. Nevertheless, 22 out of 24 SNPs in the 96% posterior probability credible set for AAD association, all in high LD with rs138300818 (D’=1, r^2^>0.89 in 1000G GBR individuals), were significant eQTLs for *THEMIS* (*p*-values < 1.9×10^−34^) and *ECHDC1* (*p*-values < 1.30×10^−8^) gene expression in whole blood, such that the minor alleles associated with protection from early onset type 1 diabetes were associated with lower *THEMIS* and *ECHCD1* expression (**Supplementary Table S8**). Our previous analysis indicated that the AAD association signal co-localises with the *THEMIS* eQTL signal (5).

### Other AAD SNPs affecting thymocyte motifs

Another SNP changing the motif-sequence similarity was rs113297984 for a motif in state 12 (active TSS) in the CD3^+^CD4^+^CD8^+^ thymocytes: rs113297984 reference (major) G allele had higher sequence motif similarity (p=9.16×10^−17^, score=47.09) than the alternative (minor) A allele (p=5.74×10^−13^, score=37.41); however, the scores were not significantly different (**Figure 4)**. While multiple TFs were predicted to bind to rs113297984 flanking sequence with DeepBind, the rs113297984 alleles did not affect their binding affinity. rs113297984 is in LD with rs138300818 (D’=1, r^2^=0.96 in 1000Genomes GBR population), and thus, has a similar eQTL profile. rs113297984 had a PCHiC interaction with *PTPRK* (CHiCAGO score 5.12) in thymus, in addition to interaction with *THEMIS* in GM12878 cell line (CHiCAGO score 11.54). Non-coding RNA overlapping the SNP was detected in CD8^+^αβ and CD3^+^CD4^+^CD8^+^ thymocytes (**Supplementary Figure S3**).

**Figure 4:**
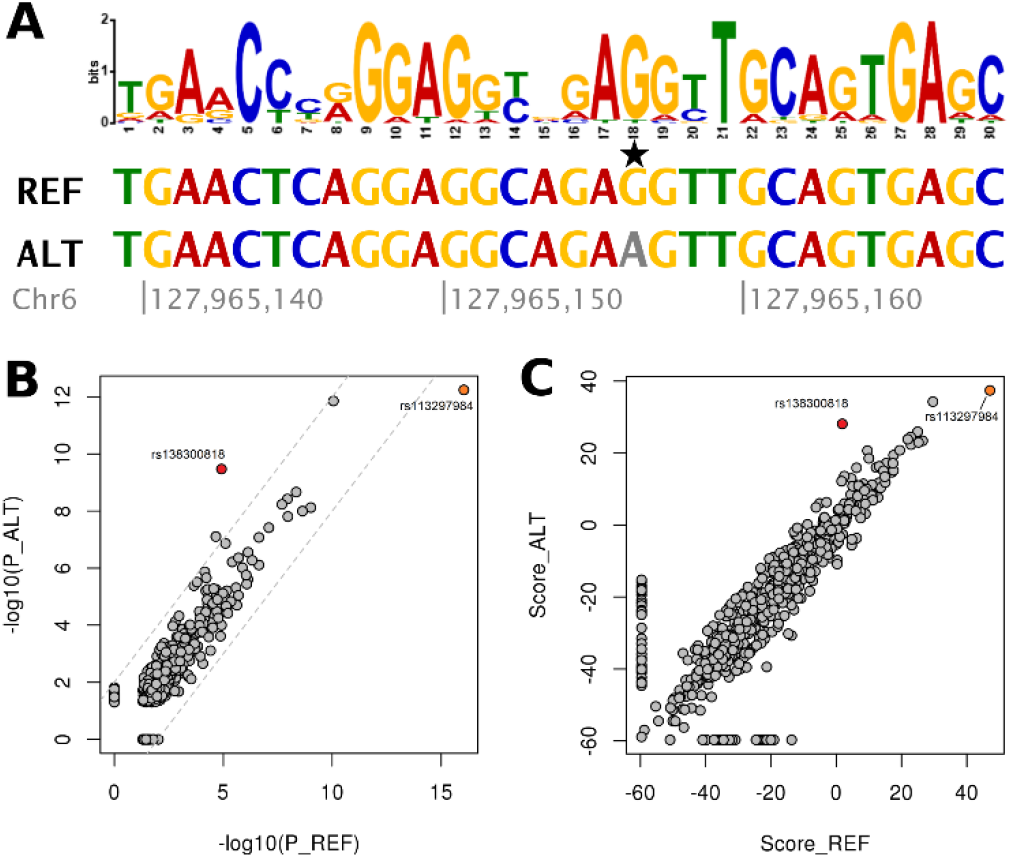
Overlap between a thymocyte motif from overlapping H3K4me3 and H3K27ac peaks (state12) in CD3^+^CD4^+^CD8^+^ double positive thymocytes and rs113297984, with significant difference in motif-sequence similarity between the reference G and alternative A alleles (highlighted with magenta background). **B** and **C:** Allelic difference in P-values (**B**) and scores in state12 peaks in CD3^+^CD4^+^CD8^+^ double positive thymocytes. rs113297984 is indicated with orange color; rs138300818 is indicated with red color for comparison.

### Type 1 diabetes SNPs affecting thymocyte motifs

We extended the same approach to other type 1 diabetes susceptibility loci. Out of 1,165 examined credible set variants from 20 susceptibility loci, 9% demonstrated allele specific sequence similarity to thymocyte motifs (E-value(p_diff_) < 5.88×10^−4^); however, only two variants had significant allelic difference in scores (E-value(Score_diff_) < 5.88×10^−4^; **Table 2**). The largest difference was observed for rs142852921 on chromosome 7p15.2, with the reference G insertion allele matching a state 10 motif (distal active promoters) in CD3^+^CD4^+^CD8^+^ cells (*p* = 4.49×10^−15^ vs. *p* = 5.04×10^−5^ for alternative allele; **Figure 5A**). Similar allele-specific motifs were observed for rs142852921 for altogether 12 thymocyte cell type – histone modification combinations. rs142852921 belongs to the finemapping block with 0.95 posterior probablility, with the reference G insertion associated with higher type 1 diabetes risk (13). We detected non-coding RNA expression in thymocyte cells overlapping rs142852921 (**Supplementary Figure S4**). rs142852921 G insertion is associated with lower *SKAP2* (whole blood *p* = 3.2×10^−6^, m-value = 1) and *HOXA7* (adipose and thyroid tissues, *p* < 4.9×10^−5^, m-value = 1) expression.

**Table 2:** Thymocyte motifs overlapping credible set SNPs for AAD, type 1 diabetes and other autoimmune diseases with marked difference in motif - sequence similarity between the reference and alternative alleles (E-value for difference in scores < 5.88×10^−4^). For each SNP, only the histone modification and cell type combination with the largest allelic difference is shown. For AAD, results are shown additionally given for rs113297984 with significant E_Diff_ (P) but non-significant E_Diff_ (Sc).

| Pheno | SNP | REF | ALT | Locus | $P_{\text{REF}}$ | $P_{\text{ALT}}$ | $S_{\text{REF}}$ | $S_{\text{ALT}}$ | $E_{\text{Diff}}(\text{P})$ | $E_{\text{Diff}}(\text{Sc})$ | HistMod | Cell type |
| --- | --- | --- | --- | --- | --- | --- | --- | --- | --- | --- | --- | --- |
| AAD | rs138300818 | - | G | 6p22.33 | $1.12 \times 10^{-4}$ | $2.53 \times 10^{-13}$ | -17.9 | 38.7 | $2.51 \times 10^{-29}$ | $6.57 \times 10^{-7}$ | H3K27ac | CD4<sup>+</sup>αβ |
| AAD | rs113297984 | G | A | 6p22.33 | $9.16 \times 10^{-17}$ | $5.74 \times 10^{-13}$ | 47.1 | 37.4 | $8.25 \times 10^{-10}$ | 0.317 | state12 | CD3<sup>+</sup>CD4<sup>+</sup>CD8<sup>+</sup> |
| T1D | rs142852921 | G | - | 7p15.2 | $4.49 \times 10^{-15}$ | $5.04 \times 10^{-5}$ | 43.5 | -11.6 | $5.4 \times 10^{-50}$ | $3.6 \times 10^{-7}$ | state10 | CD3<sup>+</sup>CD4<sup>+</sup>CD8<sup>+</sup> |
| T1D | rs66733041 | CT(ATG)<sub>2</sub>ATAC | - | *AFF3* | $1.35 \times 10^{-11}$ | 0.0067 | 31.3 | -41.2 | $3.0 \times 10^{-32}$ | $3.8 \times 10^{-4}$ | H3K4me3 | CD4<sup>+</sup>αβ |

Pheno: Original phenotype for the association. REF: Reference allele; ALT: Alternative allele. Locus: chromosomal location or a nearby gene. $P_{\text{REF}}$ and $P_{\text{ALT}}$ : P-value for similarity between the thymocyte motif and the SNP flanking sequence with the REF and ALT alleles, respectively. $S_{\text{REF}}$ and $S_{\text{ALT}}$ : Score for similarity between the thymocyte motif and the SNP flanking sequence with the REF and ALT alleles, respectively. $E_{\text{Diff}}(\text{P})$ : E-value for the difference between the $P_{\text{REF}}$ and $P_{\text{ALT}}$ . $E_{\text{Diff}}(\text{Sc})$ : E-value for the difference between the $S_{\text{REF}}$ and $S_{\text{ALT}}$ . HistMod: Histone modification.

**Figure 5:**
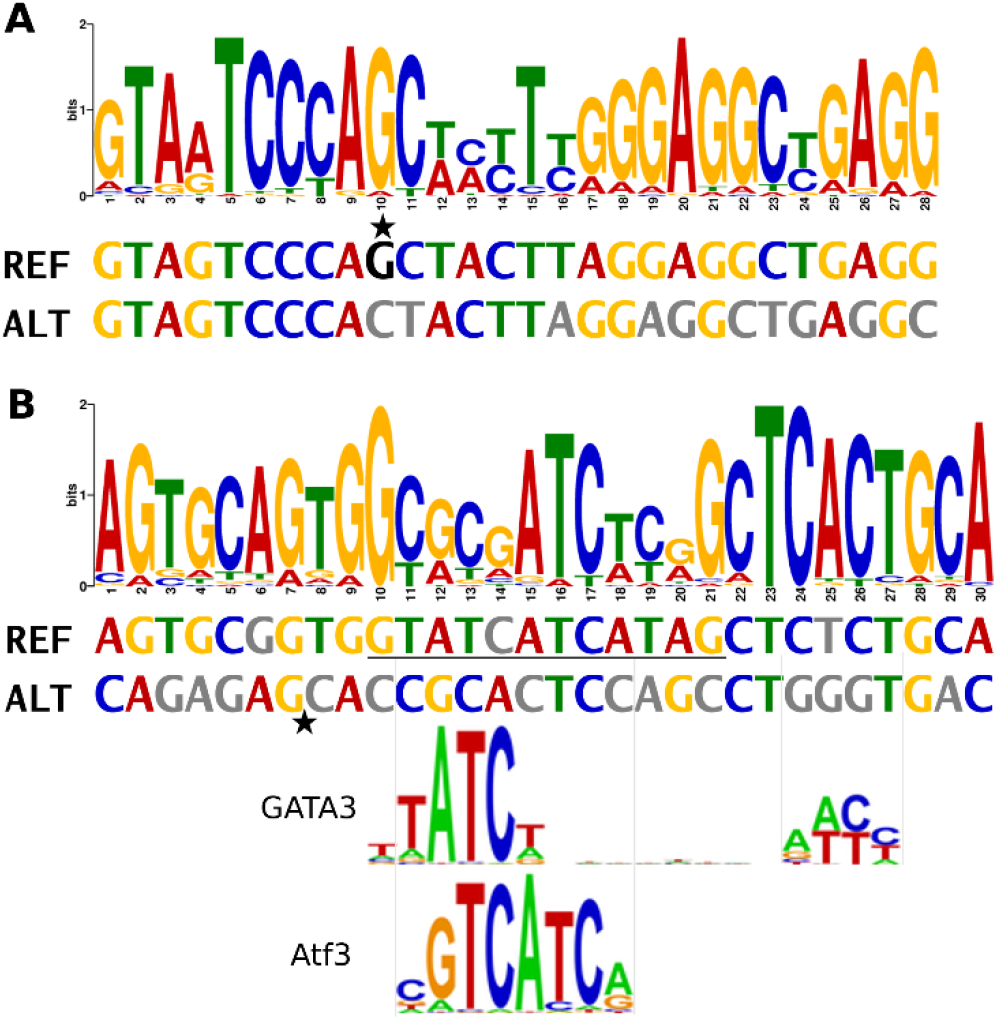
Two T1D susceptibility SNPs have allele specific thymocyte motifs. **A:** The reference G insertion allele of rs142852921 on chromosome 7p15.2 (*SKAP2*) region matches a state 10 motif (distal active promoters) in CD3^+^CD4^+^CD8^+^ cells (REF *p*=4.5×10^−15^, score 43.5; ALT p=5.0×10^−5^, score −11.6). **B**: In the *AFF3* locus, rs66733041 reference allele (underlined sequence) forms a H3K4me3 motif in CD4^+^αβ cells (p=1.4×10^−11^, score 31.1), as well as GATA3 and Aft3 binding motifs, all of which are disrupted by the 12 bp deletion allele (indicated with a star). For better match, the ALT sequence was converted to reverse complement format.

Second, the reference allele of rs66733041 on *AFF3* locus matched a thymocyte motif (**Figure 5B**, **Supplementary Table S9**) and binding sites for Atf3 and GATA3 TFs (**Supplementary Table S7**), which were all removed by the 12 bp deletion allele. However, neither of these variants had PCHiC interactions in thymocytes or in primary blood cells.

## DISCUSSION

Recent genetic analyses have reported a novel susceptibility locus associated with early diagnosis of type 1 diabetes between the *PTPRK* and *THEMIS* genes (13,30). As the thymus is likely to play a major role in early onset autoimmune diseases, and the *THEMIS* gene contributes to positive and negative T cell selection in thymus, we hypothesised that the underlying causal genetic variant would affect a regulatory motif active in thymocytes. Based on histone modification peak calls across the genome in four thymocyte cell types, we identified up to twenty motifs for each histone modification and thymocyte cell combination, representing DNA sequence motifs active in thymocytes (**Figure 1**). One of the SNPs associated with early onset type 1 diabetes in the *PTPRK-THEMIS* locus significantly affected the identified thymocyte motifs: the minor G insertion allele of rs138300818, protecting from early onset type 1 diabetes, created in multiple thymocyte cell populations a thymocyte motif, not similar to known TF-binding motifs. At the same time, the insertion allele disrupted a TF-binding motif for RFX7 and RFX5, two structurally similar RFX family members (**Figure 2**). RFX7 is abundantly expressed in thymus and lymphoid organs (31). Rfx7 coordinates a transcriptional network controlling cell metabolism in natural killer (NK) cells, and Rfx7 binding sites are found in promoter regions of genes either up or down-regulated upon RFX7 knock-out in mice (31). RFX5 regulates transcription of HLA class II genes, and mutations inactivating *RFX5* cause bare lymphocyte syndrome, a severe condition with primary HLA class II deficiency (32).

The regulatory relevance of rs138300818 was further strengthened by detection of non-coding RNA expression in all the studied thymocyte cells overlapping rs138300818. eQTL data suggested that the RFX5/7 binding rs138300818 reference allele is associated with higher *THEMIS* expression on whole blood (5). Finally, the role of rs138300818 variant in early onset type 1 diabetes was further supported by chromatin conformation interactions with both *PTPRK* and *THEMIS* in thymus and naïve CD8 cells. Whereas the role of *THEMIS* is evident in thymocytes, also *PTPRK* has been implicated in regulating development of CD4^+^ T cells (14) in addition to a variety of other cellular processes. Altogether the results suggest that the rs138300818 insertion creates a common regulatory thymocyte motif that simultaneously disrupts the RFX5/7 motif, interfering with the *THEMIS* (and *PTPRK*) gene expression, leading to protection from early onset type 1 diabetes.

We further extended the analysis to all previously fine-mapped type 1 diabetes SNPs. Only two out of the 1,165 studied type 1 diabetes associated variants significantly affected an active thymocyte motif. In *AFF3* locus, a 12 basepair insertion allele at rs66733041 created a thymocyte motif and TF binding motifs for Atf3 and GATA3. GATA3 plays an important role in T cell development, and it directly regulates many critical genes and TFs in the thymus, while demonstrating cell-type specific binding patterns and gene regulation. Variants in *GATA3* gene were recently identified as a novel susceptibility locus for type 1 diabetes (3). Furthermore, GATA3 was found to regulate active and repressive histone modifications at target enhancers, supporting the relevance of our approach (33).

All the three variants with significant allelic difference for thymocyte motif similarity (E-value(Scorediff) < 5.88×10^−4^) were indel variants of one to 12 bp long. When considering only the less stringent allelic difference in *p*-values (E-value(*P*diff) < 5.88×10^−4^), many more variants demonstrated allelic difference, including the rs113297984 G/A variant on *PTPRK-THEMIS* locus. As many disease associated non-indel variants have been suggested to alter TF binding sites (27,34), the comparative score distribution employed in this work may be too stringent and less sensitive to non-indel variants and capture only the most striking differences.

One limitation of this work is the sparsity of the thymocyte data. As only a few histone modification peaks were measured for each of the four individual, and different histone modification combinations were available for each, it was not possible to define chromatin states within each individual. This was overcome by combining histone modification peaks across different biological replicates into pseudo-chromatin states, following a previously reported annotation e.g., combining H3K4me1 and H3K27Ac signals for state 7, active enhancers (29).

It should be noted that the identified thymocyte motifs are not necessarily specific to thymocytes. This was reflected in the thymocyte motif comparison with known TF binding motifs, revealing 11 TFs, out of which most are expressed in nearly all tissues. Interestingly, SP3 has the highest expression in thymus across the FANTOM5 tissues, supporting its relevance in thymocytes. Furthermore, the thymocyte motifs affected by rs138300818 in the *PTPRK-THEMIS* were tightly linked back to thymocyte function through chromatin conformation data from thymus, and the RFX7 being highly expressed in thymus.

To conclude, the use of versatile biological data on thymocytes suggests that the rs138300818 insertion protects from early onset type 1 diabetes through affecting regulatory elements in thymocytes, including RFX5/7 binding. Functional studies are needed to confirm the finding.

## Acknowledgements

This study makes use of data generated by the Blueprint Consortium. A full list of the investigators who contributed to the generation of the data is available from www.blueprint-epigenome.eu. Funding for the project was provided by the European Union's Seventh Framework Programme (FP7/2007-2013) under grant agreement no 282510 BLUEPRINT. The Genotype-Tissue Expression (GTEx) Project was supported by the Common Fund of the Office of the Director of the National Institutes of Health, and by NCI, NHGRI, NHLBI, NIDA, NIMH, and NINDS. The data used for the analyses described in this manuscript were obtained from the GTEx Portal (https://gtexportal.org).

## Author Contributions

NS analysed the data, contributed to design and interpretation, and wrote the manuscript. ARG designed the study, contributed to data analysis and interpretation, and reviewed the manuscript. JRJI contributed to data analysis, study design and interpretation, and reviewed the manuscript. MLP, AJC and JAT contributed to study design and interpretation and reviewed the manuscript.

