## Supplementary data for "Thymocyte regulatory variant alters transcription factor binding and protects from type 1 diabetes in infants"

Supplementary Table S4: Gene expression across different tissues for the genes within 1Mbp from rs72975913 (chromosome 6, base pairs 127,293,932 - 129,293,932, GRCh38)..6

**Supplementary Table S1: Number and width of genome-wide histone modification peaks in raw data, after filtering, and in final analysis after pooling histone modification peaks from the same cell types, and after excluding exons.**

| | | | Raw peak calls | | | | Filtered peaks | | | | | | |
|---|---|---|---|---|---|---|---|---|---|---|---|---|---|
| **Histone mod** | **Cell type** | **subj** | **N** | **Width** | **Min width** | **Max width** | **RNA %** | **N** | **Width** | **Min width** | **Max width** | **N peaks final** | **N motifs** |
| H3K4me3 | $CD3^+CD4^+CD8^+$ | TH91 | 22407 | 910 | 150 | 30979 | 90 % | 18668 | 1104 | 150 | 30979 | 48295 | 17 |
| | | TH101 | 33636 | 887 | 180 | 36509 | 87 % | 25661 | 1252 | 180 | 36509 | | |
| | | TH118 | 34356 | 539 | 136 | 30024 | 87 % | 26972 | 792 | 136 | 30024 | | |
| | $CD3^-CD4^+CD8^+$ | TH89 | 18354 | 703 | 186 | 9758 | 92 % | 16125 | 787 | 186 | 9758 | 33132 | 19 |
| | | TH91 | 22947 | 722 | 130 | 13188 | 91 % | 19659 | 886 | 130 | 13188 | | |
| | $CD4^+$αß | TH91 | 26339 | 911 | 160 | 29319 | 84 % | 20448 | 1188 | 160 | 29319 | 37452 | 20 |
| | $CD8^+$αß | TH91 | 25161 | 1000 | 170 | 16113 | 85 % | 19319 | 1292 | 170 | 16113 | 36825 | 18 |
| H3K4me1 | $CD3^+CD4^+CD8^+$ | TH101 | 55622 | 1400 | 220 | 108319 | 80 % | 17551 | 2783 | 220 | 108319 | 36308 | 2 |
| H3K27ac | $CD3^+CD4^+CD8^+$ | TH91 | 14287 | 373 | 176 | 5688 | 100 % | 13029 | 395 | 176 | 5688 | 33131 | 18 |
| | | TH118 | 38531 | 250 | 140 | 6534 | 97 % | 30450 | 293 | 140 | 6534 | | |
| | $CD3^-CD4^+CD8^+$ | TH89 | 13035 | 394 | 200 | 5394 | 99 % | 11334 | 423 | 200 | 5394 | 12334 | 17 |
| | $CD4^+$αß | TH91 | 7088 | 313 | 176 | 3567 | 99 % | 6505 | 323 | 176 | 3567 | 6011 | 18 |
| H3K36me3 | $CD3^+CD4^+CD8^+$ | TH101 | 36594 | 2143 | 220 | 185670 | 100 % | 5563 | 14425 | 232 | 185670 | 53760 | 1 |
| | | TH118 | 41108 | 1333 | 190 | 135740 | 100 % | 1416 | 6843,5 | 190 | 135740 | | |
| H3K27me3 | $CD3^-CD4^+CD8^+$ | TH89 | 14696 | 1099 | 180 | 116374 | 58 % | 1240 | 1368 | 180 | 20301 | 2333 | 20 |

Raw peak calls: unmodified histone modification peak calls from Encode data; filtered peaks: peaks after filtering for those with fold change ≥3, q-value<0.0001, and some level of total RNA sequencing overlapping the peak, indicative of active chromatin region; N peaks final: number of peaks after pooling all peaks from the same cell type, and excluding exons (potentially leading to splitting a peak into multiple new peak segments). N motifs: Number of significant DNA motifs (E-value < $2.94 \times 10^{-5}$) underlying each cell type specific histone modification peak, identified with MEME software.

**Supplementary Table S2: Number of peaks for combined chromatin states**

| Chromatin state | combination of histone modification marks | Thymocyte cell type | Number of peaks | Number of motifs |
|---|---|---|---|---|
| state7 | H3K4me1 and H3K36me3 | CD3+ CD4+ CD8+ | 3524 | 9 |
| state9 | H3K4me1 and H3K27Ac | CD3+ CD4+ CD8+ | 18906 | 16 |
| state10 | H3K4me3, H3K27Ac, and H3K4me1 | CD3+ CD4+ CD8+ | 14433 | 14 |
| state11 | H3K4me3 and H3K4me1 | CD3+ CD4+ CD8+ | 28932 | 16 |
| state12 | H3K4me3 and H3K27Ac | CD3+ CD4+ CD8+ | 24341 | 15 |
| state12 | H3K4me3 and H3K27Ac | CD3- CD4+ CD8+ | 10390 | 19 |
| state12 | H3K4me3 and H3K27Ac | CD4+ αß | 5741 | 14 |

Number of motifs: Number of significant DNA motifs underlying each celltype specific composite histone state peak, identified with MEME software. E-value for statistical significance was adjusted for multiple testing, E-value<0.01/17 cell type – histone modification or state combinations/20 motifs searched for each combination=$2.94\times10^{-5}$

**Supplementary Table S3:** High-confidence chromatin 3D conformation capture interactions (CHiCAGO score > 5) in 16 primary blood cells and in foetal thymus for the 58 SNPs in the credible set of likely causal SNPs for age at diabetes diagnosis.

| Chr:pos_b37 | SNP | Finemap | Fragment | Tissue | Gene | ChiCAGO |
|---|---|---|---|---|---|---|
| 6:128265918 | rs802750 | 0.50 | | | | |
| 6:128266249 | rs6939352 | 0.96 | *chr6:128,266,169..128,271,884 (5.71KB)* | Foetal thymus | *PTPRK* | 8.18 |
| 6:128268564 | rs802747 | 0.50 | | Naive CD4 | *PTPRK* | 5.28 |
| 6:128269179 | rs802746 | 0.42 | | | | |
| 6:128270066 | rs9491889 | 0.96 | | | | |
| 6:128270122 | rs9491890 | 0.96 | | | | |
| 6:128272323 | rs802744 | 0.50 | *chr6:128,271,885..128,279,291 (7.41KB)* | Foetal thymus | *PTPRK* | 7.6 |
| 6:128272875 | rs802743 | 0.50 | | | | |
| 6:128276699 | rs802740 | 0.50 | | | | |
| 6:128277150 | rs9491891 | 0.96 | | | | |
| 6:128277209 | rs802739 | 0.50 | | | | |
| 6:128277274 | rs147626184 | 0.96 | | | | |
| 6:128277932 | rs802738 | 0.50 | | | | |
| 6:128278052 | rs802737 | 0.50 | | | | |
| 6:128278121 | rs376827043 | 0.50 | | | | |
| 6:128278229 | rs1418600 | 0.50 | | | | |
| 6:128278230 | rs1418601 | 0.50 | | | | |
| 6:128278232 | rs118097399 | 0.96 | | | | |
| 6:128278335 | rs802735 | 0.50 | | | | |
| 6:128278797 | rs802734 | 0.42 | | | | |
| 6:128279184 | rs802733 | 0.50 | | | | |
| 6:128279421 | rs802732 | 0.50 | *chr6:128,279,292…128,279,651 (0.36KB)* | | | |
| 6:128279428 | rs802731 | 0.42 | | | | |
| 6:128279496 | rs35576497 | 0.50 | | | | |
| 6:128280103 | rs802730 | 0.42 | *chr6:128,279,652..128,284,437 (4.79KB)* | Foetal thymus | *PTPRK* | 5.62 |
| 6:128280357 | rs9491892 | 0.96 | | | | |
| 6:128280374 | rs9482848 | 0.96 | | | | |
| 6:128280930 | rs9491893 | 0.96 | | | | |
| 6:128281555 | rs802728 | 0.50 | | | | |
| 6:128281660 | rs802727 | 0.50 | | | | |
| 6:128281860 | rs802726 | 0.50 | | | | |
| 6:128282028 | rs802725 | 0.42 | | | | |
| 6:128282757 | rs1089653 | 0.42 | | | | |
| 6:128282782 | rs1089652 | 0.50 | | | | |
| 6:128283192 | rs802724 | 0.50 | | | | |
| 6:128284218 | rs802722 | 0.50 | | | | |
| 6:128284770 | rs802721 | 0.50 | *chr6:128,284,438..128,288,435 (4.00KB)* | Foetal thymus | *PTPRK* | 5.12 |
| 6:128286300 | rs113297984 | 0.96 | | | | |
| 6:128286385 | rs72973797 | 0.96 | | | | |
| 6:128287157 | rs72973800 | 0.96 | | | | |
| 6:128287847 | rs761332 | 0.96 | | | | |

| | | | | | |
|---|---|---|---|---|---|
| 6:128288535 rs9482849 | 0.96 | *chr6:128,288,436..128,290,187 (1.75KB* | Foetal thymus | *PTPRK* | 11.35 |
| 6:128289018 rs802719 | 0.42 | | | | |
| 6:128289213 rs12111314 | 0.96 | | | | |
| 6:128291198 rs3190930 | 0.42 | *chr6:128,290,677..128,298,505 (7.83KB)* | Foetal thymus | *PTPRK* | 13.79 |
| 6:128291648 rs41285280 | 0.42 | | Naive CD8 | *THEMIS* | 5.65 |
| 6:128291680 rs11753289 | 0.96 | | | | |
| 6:128292391 rs4559105 | 0.42 | | | | |
| 6:128293505 rs9482850 | 0.96 | | | | |
| 6:128293561 rs55743914 | 0.42 | | | | |
| 6:128293633 rs9482851 | 0.96 | | | | |
| 6:128293931 rs72975913 | 0.96 | | | | |
| 6:128294054 rs72975916 | 0.96 | | | | |
| 6:128294708 rs35469349 | 0.42 | | | | |
| 6:128295501 rs7738609 | 0.96 | | | | |
| 6:128297021 rs138300818 | 0.96 | | | | |
| 6:128297603 rs3901020 | 0.96 | | | | |
| 6:128297610 rs4510698 | 0.96 | | | | |

Chromosome position is given in b37 coordinates. Finemap: posterior probability of causal variant within the group. Fragment: the PCHiC fragment used as the target in the experiment. Gene: the gene used as bait in the experiment. ChiCAGO: ChiCAGO score for the PCHiC interaction. Score ≥5 was considered significant. No interactions were detected in pancreatic islets (Miguel-Escalada I et al.: Human pancreatic islet 3D chromatin architecture provides insights into the genetics of type 2 diabetes. bioRxiv. :400291, 2018.)

**Supplementary Table S4: Gene expression across different tissues for the genes within 1Mbp from rs72975913 (chromosome 6, base pairs 127,293,932 - 129,293,932, GRCh38).**

| Gene | ROADMAP RNA expression | Human Protein Atlas expression | Thymus [tpm] |
|---|---|---|---|
| *RSPO3*: R-spondin 3 | | Low all/ smooth muscle, endometrium | 2 |
| *RNF146*: Ring finger protein 146 | Thymus, primary mononuclear cells, CD4 Memory Primary cells, CD4 naive primary cells, primary CD8+ naïve T cells | All<br>Blood: all | 25.9 |
| *ECHDC1*: Ethylmalonyl-CoA decarboxylase 1 | Thymus, primary mononuclear cells, CD4 Memory Primary cells, CD4 naive primary cells, primary CD8+ naïve T cells | All<br>Blood: all | 28.9 |
| *KIAA0408* | CD4 Memory Primary cells | Brain/ mixed | |
| *SOGA3*: SOGA family member 3 | CD4 Memory Primary cells | Brain<br>Blood: T-cell enriched | |
| C6orf58 | | duodenum, salivary gland, stomach | |
| *THEMIS*: Thymocyte selection associated | Thymus, primary mononuclear cells, CD4 Memory Primary cells, CD4 naive primary cells, primary CD8+ naïve T cells | **Thymus enriched**<br>**Blood: T-cell enriched** | **322.9** |
| *PTPRK*: Protein tyrosine phosphatase, receptor type K | Thymus, primary CD8+ naïve T cells | All<br>Blood: B-cell and T-cell enriched | 102.5 |

Roadmap RNA expression: RNA expression in blood cells (including CD4 Memory Primary cells, CD4 naive primary cells, primary CD8+ naïve T cells) and in thymus. **Human Protein Atlas expression**: Summary of protein and RNA expression in Human Protein Atlas in HPA, GTEx, and FANTOM5 data sets; and in the blood atlas. Thymus: RNA expression in thymus [transcripts per million, tpm] in the Fantom5 data base.

**Supplementary Table S5: Overlap of discovered thymocyte motifs with known motifs**

| Histone_cell_type | Thymocyte motif ID | Target ID | Target name | Data base | p-value | E-value | q-value |
|---|---|---|---|---|---|---|---|
| **H3K4me3_CD3n_CD4p_CD8p** | **YCTCCCTCYCYCYCYYYCTCY** | **MA0528.1** | **ZNF263** | **Jaspar core** | **3.74E-11** | **6.76E-08** | **1.35E-07** |
| state12_CD3p_CD4p_CD8p | CTTYCYCYCTCYCYYYCYCYTCCTCYYHCY | MA0528.1 | ZNF263 | Jaspar core | 1.06E-10 | 1.91E-07 | 3.82E-07 |
| **H3K4me3_CD3n_CD4p_CD8p** | **GSGGCGGBGSGGGSVGGRGGCGGGGCSGGS** | **MA0516.1** | **SP2** | **Jaspar core** | **3.69E-10** | **6.68E-07** | **1.32E-06** |
| state12_CD3n_CD4p_CD8p | GGRGGAGGRGGAGGR | MA0528.1 | ZNF263 | Jaspar core | 5.02E-10 | 9.07E-07 | 1.81E-06 |
| H3K27ac_CD3p_CD4p_CD8p | GGGGGDSGRGGMGGRGGSRGG | MA0528.1 | ZNF263 | Jaspar core | 1.67E-09 | 3.02E-06 | 6.00E-06 |
| H3K27ac_CD4_ab | RGGRRRGGGRAGGRRARRRGR | MA0528.1 | ZNF263 | Jaspar core | 2.16E-09 | 3.90E-06 | 7.79E-06 |
| state9_CD3p_CD4p_CD8p | CCCTCYYYCTCYYCC | MA0528.1 | ZNF263 | Jaspar core | 3.73E-09 | 6.74E-06 | 1.35E-05 |
| H3K4me3_CD3p_CD4p_CD8p | CCSSSCCCHSRSCCCSCCCCSSCCSCSCSS | MA0516.1 | SP2 | Jaspar core | 4.36E-09 | 7.88E-06 | 1.56E-05 |
| H3K4me3_CD3n_CD4p_CD8p | CCCSSSCCCCGCCSCCGCCCSCSSCSCSCC | MA0516.1 | SP2 | Jaspar core | 8.69E-09 | 1.57E-05 | 3.11E-05 |
| state12_CD4_ab | KGRGGRWGGGGRAGSRGRGRV | MA0528.1 | ZNF263 | Jaspar core | 1.09E-08 | 1.97E-05 | 3.94E-05 |
| H3K27ac_CD3n_CD4p_CD8p | CSCSSCCSSCCSCBCCCCCCSSCCCBSSSC | MA0516.1 | SP2 | Jaspar core | 1.54E-08 | 2.79E-05 | 5.52E-05 |
| **H3K4me3_CD3n_CD4p_CD8p** | **GSGGCGGBGSGGGSVGGRGGCGGGGCSGGS** | **MA0079.3** | **SP1** | **Jaspar core** | **4.01E-08** | **7.26E-05** | **7.19E-05** |
| H3K27ac_CD4_ab | CCCCCGCCCCSCCCCSSCCSSCSCCGCCS | MA0516.1 | SP2 | Jaspar core | 2.08E-08 | 3.76E-05 | 7.46E-05 |
| H3K27ac_CD3p_CD4p_CD8p | GGGGGDSGRGGMGGRGGSRGG | MA0516.1 | SP2 | Jaspar core | 4.20E-08 | 7.60E-05 | 7.55E-05 |
| H3K4me3_CD4_ab | CSSSSCSCSCSCCSSCCCCGCCSCSSCCS | MA0516.1 | SP2 | Jaspar core | 2.89E-08 | 5.23E-05 | 1.03E-04 |
| state11_CD3p_CD4p_CD8p | GSGSSSGGCSGGGGGSGGRGSSGGSGGSSG | MA0516.1 | SP2 | Jaspar core | 3.42E-08 | 6.19E-05 | 1.23E-04 |
| H3K27ac_CD4_ab | CSSSCSSCGCCCCSGCCSCSS | MA0516.1 | SP2 | Jaspar core | 3.65E-08 | 6.59E-05 | 1.31E-04 |
| state12_CD3n_CD4p_CD8p | GSSGGSGSGGGGSSGGGCGGGGSGCGSGSS | MA0516.1 | SP2 | Jaspar core | 4.22E-08 | 7.64E-05 | 1.51E-04 |
| H3K4me3_CD3p_CD4p_CD8p | SGGSSSGGGSCGGGVGCGGVGSSGSSSGSG | MA0516.1 | SP2 | Jaspar core | 4.32E-08 | 7.82E-05 | 1.55E-04 |
| H3K4me3_CD8_ab | SGSSSGSSGSGGSSGSGGCNGGGSGSGGGG | MA0516.1 | SP2 | Jaspar core | 5.12E-08 | 9.26E-05 | 1.83E-04 |
| H3K4me3_CD3n_CD4p_CD8p | GSGGSGGGSGGSGSGGGCGSGGSGSGSGGS | MA0516.1 | SP2 | Jaspar core | 5.12E-08 | 9.25E-05 | 1.84E-04 |
| H3K27ac_CD3p_CD4p_CD8p | GSSSSSGSGGSGGMGSGGGSSGGGSCGGGG | MA0516.1 | SP2 | Jaspar core | 6.01E-08 | 1.09E-04 | 2.15E-04 |
| H3K4me3_CD8_ab | GGGGSSGGGGCGGGGGCGSGG | MA0516.1 | SP2 | Jaspar core | 6.04E-08 | 1.09E-04 | 2.17E-04 |
| H3K4me3_CD8_ab | GGGGGCGGGGCSGGGSGGSSSGGGSSSGSG | MA0079.3 | SP1 | Jaspar core | 9.05E-08 | 1.64E-04 | 2.23E-04 |
| H3K4me3_CD8_ab | GGGGGCGGGGCSGGGSGGSSSGGGSSSGSG | MA0516.1 | SP2 | Jaspar core | 1.24E-07 | 2.25E-04 | 2.23E-04 |
| H3K4me3_CD8_ab | GGGGGCGGSGCGSGG | MA0079.3 | SP1 | Jaspar core | 8.08E-08 | 1.46E-04 | 2.90E-04 |
| H3K4me3_CD4_ab | CSCSCCCCSCSCCGCCCCCGCCCSCSCSC | MA0516.1 | SP2 | Jaspar core | 9.84E-08 | 1.78E-04 | 3.53E-04 |
| **H3K27ac_CD3p_CD4p_CD8p** | **GGGGGDSGRGGMGGRGGSRGG** | **UP00021_1** | **Zfp281 (Znf281 )** | **uniprobe_mouse** | **3.13E-07** | **5.65E-04** | **3.75E-04** |
| state12_CD4_ab | CSCCGCSSSCSCCSSCSCCSCCSSSSCCGC | MA0516.1 | SP2 | Jaspar core | 1.36E-07 | 2.46E-04 | 4.86E-04 |
| H3K27ac_CD4_ab | CCCCCGCCCCSCCCCSSCCSSCSCCGCCS | UP00021_1 | Zfp281 (Znf281 ) | uniprobe_mouse | 2.75E-07 | 4.97E-04 | 4.93E-04 |
| H3K4me3_CD3n_CD4p_CD8p | CCCSSSCCCCGCCSCCGCCCSCSSCSCSCC | MA0079.3 | SP1 | Jaspar core | 2.84E-07 | 5.14E-04 | 5.09E-04 |
| **H3K4me3_CD3n_CD4p_CD8p** | **GSGGCGGBGSGGGSVGGRGGCGGGGCSGGS** | **UP00002_1** | **Sp4** | **uniprobe_mouse** | **4.49E-07** | **8.12E-04** | **5.36E-04** |
| state12_CD3p_CD4p_CD8p | SCSSCSSCGCCSSCCCCGCCCCSSCCCSC | MA0516.1 | SP2 | Jaspar core | 1.58E-07 | 2.86E-04 | 5.66E-04 |

| Histone_cell_type | Thymocyte motif ID | Target ID | Target name | Data base | p-value | E-value | q-value |
|---|---|---|---|---|---|---|---|
| H3K4me3_CD8_ab | YCTCYCTCYCYCTCY | MA0528.1 | ZNF263 | Jaspar core | 1.73E-07 | 3.12E-04 | 6.25E-04 |
| H3K27ac_CD4_ab | CSSSCSSCGCCCCSGCCSCSS | MA0079.3 | SP1 | Jaspar core | 4.15E-07 | 7.50E-04 | 7.42E-04 |
| H3K4me3_CD4_ab | CSSCGCCCCCKCCCCSGCSCBSCSCCBCCS | MA0516.1 | SP2 | Jaspar core | 2.22E-07 | 4.02E-04 | 7.96E-04 |
| state12_CD3p_CD4p_CD8p | GGGGSGSSCGGGGCGGSGSVGGSSSCGGG | MA0516.1 | SP2 | Jaspar core | 2.33E-07 | 4.21E-04 | 8.34E-04 |
| state11_CD3p_CD4p_CD8p | CSCSSSCCCSSSCSCSCSCCSSCCSCGCSC | MA0516.1 | SP2 | Jaspar core | 2.40E-07 | 4.33E-04 | 8.59E-04 |
| state12_CD3n_CD4p_CD8p | SCSSSCSSCGCCSCCSSCSCC | MA0516.1 | SP2 | Jaspar core | 2.75E-07 | 4.96E-04 | 9.75E-04 |
| **H3K4me3_CD8_ab** | **GGGGGCGGSGCGSGG** | **KLF16_DBD** | **KLF16** | **Jolma2013** | **8.17E-07** | **1.48E-03** | **9.77E-04** |
| H3K4me3_CD8_ab | GGGGGCGGSGCGSGG | MA0741.1 | KLF16 | Jaspar core | 8.17E-07 | 1.48E-03 | 9.77E-04 |
| state12_CD4_ab | CCSSCSSCSCCCCSGCCSCCBCCCKSSCC | MA0516.1 | SP2 | Jaspar core | 2.84E-07 | 5.14E-04 | 1.02E-03 |
| state9_CD3p_CD4p_CD8p | SSGGSSSGGSCGGGGSCGGGGGBSGSSGG | MA0516.1 | SP2 | Jaspar core | 2.94E-07 | 5.32E-04 | 1.05E-03 |
| H3K4me3_CD8_ab | GGGGGCGGGGCSGGGSGGSSSGGGSSSGSG | UP00021_1 | Zfp281 (Znf281 ) | uniprobe_mouse | 9.61E-07 | 1.74E-03 | 1.15E-03 |
| H3K27ac_CD4_ab | CCTCCTCYYCC | MA0528.1 | ZNF263 | Jaspar core | 3.39E-07 | 6.13E-04 | 1.23E-03 |
| H3K4me3_CD8_ab | GGGGGCGGSGCGSGG | MA0516.1 | SP2 | Jaspar core | 1.42E-06 | 2.57E-03 | 1.27E-03 |
| H3K27ac_CD3n_CD4p_CD8p | GGSGSSSGGGGSSSSGGMSGSGGSCGGSSG | MA0516.1 | SP2 | Jaspar core | 3.58E-07 | 6.48E-04 | 1.29E-03 |
| state12_CD3p_CD4p_CD8p | GGGGAWGGRGGAGGG | MA0528.1 | ZNF263 | Jaspar core | 4.68E-07 | 8.47E-04 | 1.69E-03 |
| H3K4me3_CD3n_CD4p_CD8p | GSGGSGGGSGGSGSGGGCGSGGSGSGSGGS | MA0079.3 | SP1 | Jaspar core | 1.43E-06 | 2.58E-03 | 1.73E-03 |
| H3K4me3_CD3n_CD4p_CD8p | GSGGSGGGSGGSGSGGGCGSGGSGSGSGGS | UP00021_1 | Zfp281 (Znf281 ) | uniprobe_mouse | 1.44E-06 | 2.61E-03 | 1.73E-03 |
| H3K4me3_CD3p_CD4p_CD8p | CSCSCGCCCCSGCCCCGCCCC | MA0516.1 | SP2 | Jaspar core | 5.13E-07 | 9.28E-04 | 1.84E-03 |
| state12_CD3n_CD4p_CD8p | GSSGGSGSGGGGSSGGGCGGGGSGCGSGSS | UP00021_1 | Zfp281 (Znf281 ) | uniprobe_mouse | 1.12E-06 | 2.02E-03 | 2.00E-03 |
| H3K27ac_CD4_ab | CCCCCGCCCCSCCCCSSCCSSCSCCGCCS | MA0079.3 | SP1 | Jaspar core | 1.75E-06 | 3.16E-03 | 2.09E-03 |
| H3K27ac_CD3n_CD4p_CD8p | CSCSSCCSSCCSCBCCCCCCSSCCCBSSSC | MA0079.3 | SP1 | Jaspar core | 1.23E-06 | 2.22E-03 | 2.20E-03 |
| H3K4me3_CD3p_CD4p_CD8p | CCSSSCCCHSRSCCCSCCCCSSCCSCSCSS | MA0079.3 | SP1 | Jaspar core | 1.49E-06 | 2.69E-03 | 2.26E-03 |
| H3K4me3_CD3p_CD4p_CD8p | CCSSSCCCHSRSCCCSCCCCSSCCSCSCSS | UP00021_1 | Zfp281 (Znf281 ) | uniprobe_mouse | 1.89E-06 | 3.41E-03 | 2.26E-03 |
| H3K4me3_CD3n_CD4p_CD8p | GSGGCGGBGSGGGSVGGRGGCGGGGCSGGS | UP00021_1 | Zfp281 (Znf281 ) | uniprobe_mouse | 2.56E-06 | 4.62E-03 | 2.29E-03 |
| H3K4me3_CD3n_CD4p_CD8p | GSGGCGGBGSGGGSVGGRGGCGGGGCSGGS | KLF16_DBD | KLF16 | Jolma2013 | 4.01E-06 | 7.25E-03 | 2.39E-03 |
| H3K4me3_CD3n_CD4p_CD8p | GSGGCGGBGSGGGSVGGRGGCGGGGCSGGS | MA0741.1 | KLF16 | Jaspar core | 4.01E-06 | 7.25E-03 | 2.39E-03 |
| state11_CD3p_CD4p_CD8p | CSCSSSCCCSSSCSCSCSCCSSCCSCGCSC | UP00021_1 | Zfp281 (Znf281 ) | uniprobe_mouse | 1.35E-06 | 2.44E-03 | 2.42E-03 |
| **H3K4me3_CD8_ab** | **GGGGGCGGSGCGSGG** | **MA0599.1** | **KLF5** | **Jaspar core** | **3.39E-06** | **6.13E-03** | **2.43E-03** |
| state12_CD4_ab | CTYTYYCTCYTCCTY | MA0528.1 | ZNF263 | Jaspar core | 6.78E-07 | 1.23E-03 | 2.45E-03 |
| H3K4me3_CD8_ab | GGGGGCGGSGCGSGG | SP1_DBD | SP1 | Jolma2013 | 4.17E-06 | 7.55E-03 | 2.49E-03 |
| H3K27ac_CD3p_CD4p_CD8p | SCSSSSCCSSCSCCSCCSCCGCSGCCSCS | MA0516.1 | SP2 | Jaspar core | 7.30E-07 | 1.32E-03 | 2.61E-03 |
| state12_CD3p_CD4p_CD8p | GGGGSGSSCGGGGCGGSGSVGGSSSCGGG | MA0079.3 | SP1 | Jaspar core | 1.51E-06 | 2.72E-03 | 2.70E-03 |
| H3K27ac_CD4_ab | RGGRRRGGGRAGGRRARRRGR | UP00021_1 | Zfp281 (Znf281 ) | uniprobe_mouse | 1.51E-06 | 2.72E-03 | 2.72E-03 |
| H3K4me3_CD3n_CD4p_CD8p | CCCSSSCCCCGCCSCCGCCCSCSSCSCSCC | KLF16_DBD | KLF16 | Jolma2013 | 3.70E-06 | 6.69E-03 | 2.85E-03 |
| H3K4me3_CD3n_CD4p_CD8p | CCCSSSCCCCGCCSCCGCCCSCSSCSCSCC | MA0741.1 | KLF16 | Jaspar core | 3.70E-06 | 6.69E-03 | 2.85E-03 |

| Histone_cell_type | Thymocyte motif ID | Target ID | Target name | Data base | p-value | E-value | q-value |
|---|---|---|---|---|---|---|---|
| H3K4me3_CD3n_CD4p_CD8p | CCCSSSCCCCGCCSCCGCCCSCSSCSCSCC | UP00002_1 | Sp4 | uniprobe_mouse | 4.78E-06 | 8.65E-03 | 2.85E-03 |
| H3K4me3_CD3n_CD4p_CD8p | CCCSSSCCCCGCCSCCGCCCSCSSCSCSCC | UP00021_1 | Zfp281 (Znf281 ) | uniprobe_mouse | 4.08E-06 | 7.38E-03 | 2.85E-03 |
| H3K4me3_CD4_ab | CSSCGCCCCCKCCCCSGCSCBSCSCCBCCS | MA0079.3 | SP1 | Jaspar core | 1.61E-06 | 2.90E-03 | 2.88E-03 |
| H3K27ac_CD3n_CD4p_CD8p | GGGSSGGGSSGGGSGSGGCGGSGGCSGCGG | MA0516.1 | SP2 | Jaspar core | 8.10E-07 | 1.46E-03 | 2.90E-03 |
| H3K4me3_CD4_ab | CSSCGCSCSCSGCCCCSSCSC | MA0516.1 | SP2 | Jaspar core | 8.27E-07 | 1.49E-03 | 2.94E-03 |
| state12_CD3n_CD4p_CD8p | SCSSSCSSCGCCSCCSSCSCC | MA0079.3 | SP1 | Jaspar core | 1.71E-06 | 3.09E-03 | 3.04E-03 |
| **H3K4me3_CD8_ab** | **GGGGGCGGSGCGSGG** | **MA0746.1** | **SP3** | **Jaspar core** | **6.78E-06** | **1.23E-02** | **3.04E-03** |
| H3K4me3_CD8_ab | GGGGGCGGSGCGSGG | SP3_DBD | SP3 | Jolma2013 | 6.78E-06 | 1.23E-02 | 3.04E-03 |
| H3K4me3_CD4_ab | CSCSCCCCSCSCCGCCCCCGCCCSCSCSC | MA0079.3 | SP1 | Jaspar core | 1.73E-06 | 3.12E-03 | 3.09E-03 |
| state12_CD3p_CD4p_CD8p | GGGGAWGGRGGAGGG | UP00002_1 | Sp4 | uniprobe_mouse | 1.91E-06 | 3.46E-03 | 3.45E-03 |
| H3K4me3_CD3p_CD4p_CD8p | SGGSSSGGGSCGGGVGCGGVGSSGSSSGSG | UP00021_1 | Zfp281 (Znf281 ) | uniprobe_mouse | 1.96E-06 | 3.55E-03 | 3.51E-03 |
| H3K27ac_CD3p_CD4p_CD8p | GGGGGDSGRGGMGGRGGSRGG | MA0079.3 | SP1 | Jaspar core | 4.22E-06 | 7.64E-03 | 3.80E-03 |
| **state12_CD3n_CD4p_CD8p** | **CCCCCRYCCCCNCCCMCACCC** | **MA0073.1** | **RREB1** | **Jaspar core** | **1.07E-06** | **1.93E-03** | **3.85E-03** |
| **H3K4me3_CD8_ab** | **SGSSSGSSGSGGSSGSGGCNGGGSGSGGGG** | **MA0146.2** | **Zfx** | **Jaspar core Mouse** | **2.34E-06** | **4.23E-03** | **4.17E-03** |
| H3K4me3_CD3p_CD4p_CD8p | GSSSGGGGCGGSGGCGSSGGS | MA0516.1 | SP2 | Jaspar core | 1.20E-06 | 2.17E-03 | 4.29E-03 |
| state12_CD3n_CD4p_CD8p | GSSGGSGSGGGGSSGGGCGGGGSGCGSGSS | MA0079.3 | SP1 | Jaspar core | 3.64E-06 | 6.59E-03 | 4.35E-03 |
| H3K4me3_CD4_ab | CSSSSCSCSCSCCSSCCCCGCCSCSSCCS | MA0079.3 | SP1 | Jaspar core | 2.58E-06 | 4.66E-03 | 4.61E-03 |
| state11_CD3p_CD4p_CD8p | GSGSSSGGCSGGGGGSGGRGSSGGSGGSSG | MA0079.3 | SP1 | Jaspar core | 3.90E-06 | 7.05E-03 | 4.66E-03 |
| state11_CD3p_CD4p_CD8p | GSGSSSGGCSGGGGGSGGRGSSGGSGGSSG | UP00021_1 | Zfp281 (Znf281 ) | uniprobe_mouse | 2.73E-06 | 4.93E-03 | 4.66E-03 |
| H3K27me3_CD3n_CD4p_CD8p | CGCSGCCSCCGCCSCCSCCSC | MA0516.1 | SP2 | Jaspar core | 1.33E-06 | 2.40E-03 | 4.76E-03 |
| state12_CD4_ab | CCSSCSSCSCCCCSGCCSCCBCCCKSSCC | UP00021_1 | Zfp281 (Znf281 ) | uniprobe_mouse | 2.82E-06 | 5.09E-03 | 5.05E-03 |
| H3K4me3_CD3n_CD4p_CD8p | GSGGCGGBGSGGGSVGGRGGCGGGGCSGGS | MA0599.1 | KLF5 | Jaspar core | 1.25E-05 | 2.26E-02 | 5.10E-03 |
| H3K4me3_CD3n_CD4p_CD8p | GSGGCGGBGSGGGSVGGRGGCGGGGCSGGS | SP1_DBD | SP1 | Jolma2013 | 1.16E-05 | 2.11E-02 | 5.10E-03 |
| H3K4me3_CD3n_CD4p_CD8p | GSGGCGGBGSGGGSVGGRGGCGGGGCSGGS | MA0146.2 | Zfx | Jaspar core Mouse | 1.28E-05 | 2.32E-02 | 5.10E-03 |
| H3K4me3_CD3n_CD4p_CD8p | GSGGCGGBGSGGGSVGGRGGCGGGGCSGGS | MA0746.1 | SP3 | Jaspar core | 1.57E-05 | 2.84E-02 | 5.12E-03 |
| H3K4me3_CD3n_CD4p_CD8p | GSGGCGGBGSGGGSVGGRGGCGGGGCSGGS | SP3_DBD | SP3 | Jolma2013 | 1.57E-05 | 2.84E-02 | 5.12E-03 |
| H3K4me3_CD3n_CD4p_CD8p | GSGGCGGBGSGGGSVGGRGGCGGGGCSGGS | MA0528.1 | ZNF263 | Jaspar core | 1.72E-05 | 3.11E-02 | 5.14E-03 |
| H3K4me3_CD3n_CD4p_CD8p | CCCSSSCCCCGCCSCCGCCCSCSSCSCSCC | MA0746.1 | SP3 | Jaspar core | 1.22E-05 | 2.21E-02 | 5.47E-03 |
| H3K4me3_CD3n_CD4p_CD8p | CCCSSSCCCCGCCSCCGCCCSCSSCSCSCC | SP3_DBD | SP3 | Jolma2013 | 1.22E-05 | 2.21E-02 | 5.47E-03 |
| H3K4me3_CD3p_CD4p_CD8p | SGGSSSGGGSCGGGVGCGGVGSSGSSSGSG | MA0079.3 | SP1 | Jaspar core | 4.60E-06 | 8.31E-03 | 5.49E-03 |
| H3K4me3_CD3n_CD4p_CD8p | YCTCCCTCYCYCYCYYYCTCY | UP00021_1 | Zfp281 (Znf281 ) | uniprobe_mouse | 3.11E-06 | 5.62E-03 | 5.62E-03 |
| state10_CD3p_CD4p_CD8p | CCGCSCSCSCCSSSSCSCCCSGCCCCSSSC | MA0516.1 | SP2 | Jaspar core | 1.58E-06 | 2.85E-03 | 5.65E-03 |
| H3K4me3_CD4_ab | CSSCGCCCCCKCCCCSGCSCBSCSCCBCCS | UP00021_1 | Zfp281 (Znf281 ) | uniprobe_mouse | 4.75E-06 | 8.59E-03 | 5.68E-03 |
| H3K27ac_CD3p_CD4p_CD8p | GSSSSSGSGGSGGMGSGGGSSGGGSCGGGG | MA0079.3 | SP1 | Jaspar core | 4.79E-06 | 8.65E-03 | 5.73E-03 |
| H3K27ac_CD3p_CD4p_CD8p | GSSSSSGSGGSGGMGSGGGSSGGGSCGGGG | MA0146.2 | Zfx | Jaspar core Mouse | 4.81E-06 | 8.70E-03 | 5.73E-03 |

| Histone_cell_type | Thymocyte motif ID | Target ID | Target name | Data base | p-value | E-value | q-value |
|---|---|---|---|---|---|---|---|
| state12_CD3p_CD4p_CD8p | SCSSCSSCGCCSSCCCCGCCCCSSCCCSC | MA0079.3 | SP1 | Jaspar core | 3.31E-06 | 5.99E-03 | 5.93E-03 |
| H3K27ac_CD3p_CD4p_CD8p | SCSSSSCCSSCSCCSCCSCCGCSGCCSCS | UP00021_1 | Zfp281 (Znf281 ) | uniprobe_mouse | 3.54E-06 | 6.40E-03 | 6.34E-03 |
| state12_CD3n_CD4p_CD8p | CCCCCRYCCCCNCCCMCACCC | MA0516.1 | SP2 | Jaspar core | 5.30E-06 | 9.59E-03 | 6.36E-03 |
| state12_CD3n_CD4p_CD8p | CCCCCRYCCCCNCCCMCACCC | UP00021_1 | Zfp281 (Znf281 ) | uniprobe_mouse | 5.30E-06 | 9.59E-03 | 6.36E-03 |
| H3K27ac_CD3p_CD4p_CD8p | GSSSSSGSGGSGGMGSGGGSSGGGSCGGGG | UP00021_1 | Zfp281 (Znf281 ) | uniprobe_mouse | 9.27E-06 | 1.68E-02 | 6.63E-03 |
| H3K27ac_CD3p_CD4p_CD8p | GAGRGRARGGRRRARRGARAA | MA0528.1 | ZNF263 | Jaspar core | 1.85E-06 | 3.34E-03 | 6.68E-03 |
| H3K4me3_CD3n_CD4p_CD8p | CCCSSSCCCCGCCSCCGCCCSCSSCSCSCC | MA0599.1 | KLF5 | Jaspar core | 1.72E-05 | 3.11E-02 | 6.85E-03 |
| H3K4me3_CD8_ab | GGGGGCGGGGCSGGGSGGSSSGGGSSSGSG | KLF16_DBD | KLF16 | Jolma2013 | 1.15E-05 | 2.08E-02 | 6.85E-03 |
| H3K4me3_CD8_ab | GGGGGCGGGGCSGGGSGGSSSGGGSSSGSG | MA0741.1 | KLF16 | Jaspar core | 1.15E-05 | 2.08E-02 | 6.85E-03 |
| H3K4me3_CD8_ab | GGGGGCGGGGCSGGGSGGSSSGGGSSSGSG | MA0528.1 | ZNF263 | Jaspar core | 1.02E-05 | 1.84E-02 | 6.85E-03 |
| H3K27ac_CD3n_CD4p_CD8p | GGSGSSSGGGGSSSSGGMSGSGGSCGGSSG | UP00021_1 | Zfp281 (Znf281 ) | uniprobe_mouse | 3.92E-06 | 7.09E-03 | 7.04E-03 |
| H3K4me3_CD8_ab | GGGGSSGGGGCGGGGGCGSGG | MA0079.3 | SP1 | Jaspar core | 3.95E-06 | 7.15E-03 | 7.08E-03 |
| H3K4me3_CD4_ab | TCCTCCTCCYC | MA0528.1 | ZNF263 | Jaspar core | 1.98E-06 | 3.58E-03 | 7.16E-03 |
| H3K4me3_CD3p_CD4p_CD8p | GSSSGGGGCGGSGGCGSSGGS | MA0079.3 | SP1 | Jaspar core | 4.28E-06 | 7.73E-03 | 7.65E-03 |
| H3K4me3_CD3p_CD4p_CD8p | CSCSCGCCCCSGCCCCGCCCC | MA0599.1 | KLF5 | Jaspar core | 6.54E-06 | 1.18E-02 | 7.79E-03 |
| H3K4me3_CD3p_CD4p_CD8p | CSCSCGCCCCSGCCCCGCCCC | MA0079.3 | SP1 | Jaspar core | 4.38E-06 | 7.91E-03 | 7.79E-03 |
| H3K4me3_CD8_ab | SGSSSGSSGSGGSSGSGGCNGGGSGSGGGG | UP00021_1 | Zfp281 (Znf281 ) | uniprobe_mouse | 8.99E-06 | 1.63E-02 | 8.02E-03 |
| H3K4me3_CD8_ab | GGGGGCGGSGCGSGG | UP00021_1 | Zfp281 (Znf281 ) | uniprobe_mouse | 2.03E-05 | 3.67E-02 | 8.09E-03 |
| state9_CD3p_CD4p_CD8p | SSGGSSSGGSCGGGGSCGGGGGBSGSSGG | MA0079.3 | SP1 | Jaspar core | 5.18E-06 | 9.36E-03 | 8.16E-03 |
| state9_CD3p_CD4p_CD8p | SSGGSSSGGSCGGGGSCGGGGGBSGSSGG | MA0146.2 | Zfx | Jaspar core Mouse | 6.84E-06 | 1.24E-02 | 8.16E-03 |
| H3K4me3_CD8_ab | GGGGSSGGGGCGGGGGCGSGG | UP00021_1 | Zfp281 (Znf281 ) | uniprobe_mouse | 6.85E-06 | 1.24E-02 | 8.19E-03 |
| H3K4me3_CD4_ab | CSCSCCCCSCSCCGCCCCCGCCCSCSCSC | UP00021_1 | Zfp281 (Znf281 ) | uniprobe_mouse | 7.04E-06 | 1.27E-02 | 8.41E-03 |
| state12_CD4_ab | CSCCGCSSSCSCCSSCSCCSCCSSSSCCGC | MA0079.3 | SP1 | Jaspar core | 6.37E-06 | 1.15E-02 | 8.85E-03 |
| state12_CD4_ab | CSCCGCSSSCSCCSSCSCCSCCSSSSCCGC | UP00021_1 | Zfp281 (Znf281 ) | uniprobe_mouse | 7.42E-06 | 1.34E-02 | 8.85E-03 |
| H3K4me3_CD8_ab | GGGGGCGGGGCSGGGSGGSSSGGGSSSGSG | MA0599.1 | KLF5 | Jaspar core | 2.16E-05 | 3.90E-02 | 9.11E-03 |
| H3K4me3_CD8_ab | GGGGGCGGGGCSGGGSGGSSSGGGSSSGSG | SP1_DBD | SP1 | Jolma2013 | 2.46E-05 | 4.46E-02 | 9.11E-03 |
| **H3K4me3_CD8_ab** | **GGGGGCGGGGCSGGGSGGSSSGGGSSSGSG** | **UP00022_1** | **Zfp740 (Znf740)** | **uniprobe_mouse** | **2.42E-05** | **4.37E-02** | **9.11E-03** |
| H3K4me3_CD8_ab | GGGGGCGGGGCSGGGSGGSSSGGGSSSGSG | MA0146.2 | Zfx | Jaspar core Mouse | 2.54E-05 | 4.60E-02 | 9.11E-03 |
| H3K27ac_CD3p_CD4p_CD8p | SGCGSGGCGGGCSSGGSCSGGGGGCSSGGG | MA0516.1 | SP2 | Jaspar core | 2.59E-06 | 4.69E-03 | 9.24E-03 |
| H3K27ac_CD3n_CD4p_CD8p | CSCSSCCSSCCSCBCCCCCCSSCCCBSSSC | UP00021_1 | Zfp281 (Znf281 ) | uniprobe_mouse | 7.80E-06 | 1.41E-02 | 9.31E-03 |
| H3K4me3_CD8_ab | SGSSSGSSGSGGSSGSGGCNGGGSGSGGGG | MA0079.3 | SP1 | Jaspar core | 1.31E-05 | 2.37E-02 | 9.37E-03 |
| H3K27ac_CD4_ab | CSSSCSSCGCCCCSGCCSCSS | UP00021_1 | Zfp281 (Znf281 ) | uniprobe_mouse | 8.13E-06 | 1.47E-02 | 9.71E-03 |

A total of 48 motifs (135 thymocyte motif – transcription factor (TF) binding motif combinations) were significantly similar with $q<0.01$. Target ID/name: Known TF ID/name. Data base: data base for the known TF motifs, human unless otherwise stated. P/E/Q-values: statistical significance for motif similarity between the thymocyte motif and known TF binding motifs. For each known TF, the thymocyte motif with lowest q-value is highlighted in bold.

**Suppelmentary table S6: Thymocyte motifs overlapping credible set SNPs with marked difference (two orders of magnitude in p-value) in motif-sequence similarity between the reference and alternative alleles.** Significant motif overlap was considered as $P_{REF}$ or $P_{ALT}$ < 0.01/253 thymocyte motifs/58 SNPs = $6.81\times10^{-7}$**.** Significant allelic difference was calculated based on observed distribution of p-value differences, separately for each of the 17 histone mark – cell type combinations and defined as $E_{Diff}$ ($P_{REF}$ - $P_{ALT}$) < 0.01/17 = $5.88\times10^{-4}$. Significant values are indicated with bold.

| SNP | matched_sequence.REF/ALT | REF | ALT | hmark | Cell type | $P_{REF}$ | $P_{ALT}$ | $Sc_{REF}$ | $Sc_{ALT}$ | $E_{Diff}$ ($P_{REF}$ - $P_{ALT}$) | $E_{Diff}$ ($Sc_{REF}$ - $Sc_{ALT}$) |
|---|---|---|---|---|---|---|---|---|---|---|---|
| rs138300818 | ATGGGGTTTCACCACGTTG-/GCCAGGCTGGTC | | G | H3K27ac | CD4_ab | 0.000112 | ***2.53E-13*** | -17.9 | 38.7 | ***2.51E-29*** | ***6.57E-07*** |
| rs138300818 | GGGGTTTCACCACGTTG-/GCCGGCTGGTCT | | G | state12 | CD4_ab | 4.73E-05 | ***2.26E-12*** | -10.4 | 35.5 | ***2.89E-25*** | ***4.68E-05*** |
| rs138300818 | AAGACCAGCCTGG-/CCAACGTGGTGAAACCCC | | G | H3K27ac | CD3n_CD4p_CD8p | 1.41E-05 | ***1.32E-13*** | -9.0 | 39.5 | ***1.56E-24*** | 1.79E-02 |
| rs138300818 | GGAGTTCAAGACCAGCCTGG-/CCAACG | | G | state9 | CD3p_CD4p_CD8p | ***1.65E-08*** | ***5.58E-15*** | 17.0 | 43.4 | ***6.32E-23*** | 6.75E-02 |
| rs138300818 | GTTTCACCACGTTG-/GCCGGCTGGTCTTG | | G | H3K4me3 | CD3n_CD4p_CD8p | 0.000333 | ***4.94E-11*** | -15.6 | 30.7 | ***1.59E-20*** | 1.65E-02 |
| rs138300818 | ACCACGTTG-/GCCGGCTGGTCT | | G | H3K4me3 | CD3p_CD4p_CD8p | 4.86E-05 | ***6.49E-12*** | -9.0 | 34.4 | ***2.77E-20*** | 5.49E-03 |
| rs138300818 | CAGGAGTTCAAGACCAGCCTGG-/CCAACGTG | | G | H3K27me3 | CD3n_CD4p_CD8p | ***2E-09*** | ***7.88E-16*** | 20.8 | 45.9 | ***5.51E-18*** | 6.70E-02 |
| rs138300818 | ACCAGCCTGG-/CCAACGTGGTGAAACCCCATC | | G | H3K4me3 | CD4_ab | 0.000515 | ***1.34E-09*** | -15.7 | 25.7 | ***4.88E-14*** | 3.09E-02 |
| rs138300818 | CCACGTTG-/GCCGGCTGGTCTTGAAC | | G | state12 | CD3p_CD4p_CD8p | 1.21E-05 | ***3.39E-10*** | 1.8 | 28.1 | ***7.47E-13*** | 1.09E-02 |
| rs138300818 | CACGTTG-/GCCGGCTGGTCTTGAACTC | | G | state10 | CD3p_CD4p_CD8p | 2.69E-06 | ***1.35E-10*** | 7.6 | 29.7 | ***1.88E-12*** | 1.51E-02 |
| rs138300818 | CAGGAGTTCAAGACCAGCCTGG-/CCAACGTGG | | G | H3K4me3 | CD8_ab | ***9.23E-11*** | ***8.86E-16*** | 26.2 | 46.4 | ***3.37E-12*** | 1.41E-01 |
| rs138300818 | AGAGATGGGGTTTCACCACGTTG-/GCCGGC | | G | state9 | CD3p_CD4p_CD8p | 4.08E-06 | ***2.62E-10*** | 1.1 | 27.3 | ***1.98E-10*** | 6.96E-02 |
| rs138300818 | CCAGCCTGG-/CCAACGT | | G | H3K4me3 | CD4_ab | 0.00151 | ***4.10E-08*** | -4.9 | 21.6 | ***7.95E-10*** | 1.67E-01 |
| rs138300818 | GGTCAGGAGTTCAAGACCAGCCTGG-/CCAAC | | G | state7 | CD3p_CD4p_CD8p | ***3.14E-14*** | ***8.84E-18*** | 37.6 | 54.1 | ***2.42E-09*** | 2.04E-01 |
| rs138300818 | TCAAGACCAGCCTGG-/CCAACGT | | G | H3K27me3 | CD3n_CD4p_CD8p | 7.73E-06 | ***7.24E-09*** | -7.6 | 19.0 | ***4.99E-05*** | 5.28E-02 |
| rs138300818 | CCTGG-/CCAACGTGGTGAAACCCCATCTCTAC | | G | state12 | CD3n_CD4p_CD8p | ***1.64E-08*** | ***4.94E-11*** | 22.4 | 30.3 | ***1.42E-04*** | 4.93E-01 |
| rs113297984 | TGAACTCAGGAGGCAGAG/AGTTGCAGTGAGC | G | A | state12 | CD3p_CD4p_CD8p | ***9.16E-17*** | ***5.74E-13*** | 47.1 | 37.4 | ***8.25E-10*** | 3.07E-01 |
| rs1089652 | C/TTTATTTTTTT | C | T | state12 | CD3n_CD4p_CD8p | 0.000026 | ***4.19E-08*** | 10.3 | 21.5 | ***2.43E-05*** | 3.26E-01 |
| rs1089652 | GAAAAAAATAAG/ATAT | C | T | state12 | CD3n_CD4p_CD8p | 2.88E-05 | ***7.47E-08*** | 9.8 | 20.7 | ***9.44E-05*** | 3.35E-01 |
| rs1089652 | ATAC/TTTATTTTTTTCTCTTAATTTTTGTCT | C | T | state12 | CD4_ab | ***7.91E-10*** | ***1.40E-12*** | 26.7 | 34.3 | ***1.22E-04*** | 5.53E-01 |
| rs1089652 | AGAGAAAAAAATAAG/A | C | T | state12 | CD3p_CD4p_CD8p | 0.000022 | ***7.86E-08*** | 9.5 | 20.7 | ***1.37E-04*** | 2.96E-01 |
| rs1089652 | AAAAATTAAGAGAAAAAAATAAG/ATATCAA | C | T | H3K27ac | CD3p_CD4p_CD8p | ***2.06E-07*** | ***1.10E-09*** | 18.7 | 26.0 | 2.42E-03 | 5.24E-01 |
| rs1089652 | AAAAAAATAAG/A | C | T | H3K27me3 | CD3n_CD4p_CD8p | 8.85E-06 | ***6.68E-08*** | 10.3 | 21.5 | 4.73E-03 | 4.27E-01 |
| rs1089652 | C/TTTATTTTTTTCTCT | C | T | H3K27ac | CD3n_CD4p_CD8p | 2.42E-05 | ***1.61E-07*** | 8.8 | 19.9 | 7.63E-03 | 6.47E-01 |
| rs72973797 | AAATAAAATAAAATAAAG/ATAAATATAAAA | G | A | H3K27ac | CD3n_CD4p_CD8p | ***6.38E-09*** | ***3.03E-12*** | 23.8 | 30.8 | ***3.35E-05*** | 8.04E-01 |
| rs72973797 | AAATAAAATAAAG/ATA | G | A | state12 | CD3n_CD4p_CD8p | 1.36E-05 | ***2.58E-08*** | 11.7 | 21.9 | ***3.92E-05*** | 3.74E-01 |
| rs72973797 | ATATTTAC/TTTTATTTTATTTTATTTCTTGA | G | A | H3K4me3 | CD3n_CD4p_CD8p | ***3.61E-11*** | ***3.42E-14*** | 28.9 | 33.6 | ***4.97E-05*** | 8.53E-01 |
| rs72973797 | AAAATAAAG/ATAAATATAAAAACAAGAAAAG | G | A | H3K4me3 | CD3p_CD4p_CD8p | ***6.97E-10*** | ***1.95E-12*** | 26.9 | 34.9 | 6.80E-04 | 6.37E-01 |
| rs72973797 | AAAG/ATAAATATAAAA | G | A | state12 | CD4_ab | ***2.18E-07*** | ***1.03E-09*** | 19.1 | 24.7 | 1.23E-03 | 6.76E-01 |

| | | | | | | | | | | | |
|---|---|---|---|---|---|---|---|---|---|---|---|
| rs72973797 | AAAG/ATAAATATAAAA | G | A | state11 | CD3p_CD4p_CD8p | 2.66E-06 | ***1.17E-08*** | 14.5 | 23.2 | 1.76E-03 | 7.11E-01 |
| rs72973797 | TTTTTATATTTAC/TTTTATTTT | G | A | state11 | CD3p_CD4p_CD8p | ***7.53E-08*** | ***4.48E-10*** | 16.6 | 27.7 | 3.21E-03 | 6.22E-01 |
| rs72973797 | AAATAAAATAAAG/ATAAATATA | G | A | H3K4me3 | CD3n_CD4p_CD8p | ***2.71E-08*** | ***1.97E-10*** | 21.3 | 29.1 | 4.50E-03 | 7.26E-01 |
| rs72973797 | AAAATAAAG/ATAAATA | G | A | H3K27ac | CD3n_CD4p_CD8p | 8.73E-07 | ***4.87E-09*** | 16.5 | 24.6 | 5.66E-03 | 7.62E-01 |
| rs11753289 | TAAACAGATAAAAAATTA/CTAGATATTTAAA | T | G | H3K27ac | CD3n_CD4p_CD8p | ***1.51E-10*** | ***1.34E-07*** | 29.1 | 18.3 | ***7.83E-05*** | 5.07E-01 |
| rs11753289 | AAAAATTA/CTAGATATTTAAAA | T | G | H3K27ac | CD3p_CD4p_CD8p | ***8.53E-08*** | 4.45E-05 | 20.3 | 9.6 | ***2.92E-04*** | 3.77E-01 |
| rs11753289 | ATATTTTCTATCAATTTTTTAAATATCTAT/G | T | G | H3K4me3 | CD4_ab | ***4.76E-11*** | ***1.31E-08*** | 29.4 | 22.8 | 6.53E-04 | 7.20E-01 |
| rs41285280 | AAAATATGCG/AGCAAAGATAAAATGTCTTT | C | T | state12 | CD3n_CD4p_CD8p | 0.000064 | ***4.46E-07*** | 8.6 | 17.9 | 1.16E-03 | 4.15E-01 |
| rs802733 | ATTTTTTATTTGAGTTCAGTATTGGTCG/ATA | G | A | H3K4me3 | CD4_ab | 2.73E-05 | ***1.94E-07*** | 9.1 | 18.8 | 4.39E-03 | 6.19E-01 |
| rs12111314 | TAACATAATAAACATTACC/TGGGTAAAAAA | C | T | H3K27ac | CD4_ab | 2.85E-05 | ***2.04E-07*** | 10.3 | 19.0 | 6.51E-03 | 4.81E-01 |

$P_{REF}$ and $P_{ALT}$: p-value for motif overlap with the DNA sequence containing the REF/ALT alleles. $Sc_{REF}$ and $Sc_{ALT}$: Score value for motif overlap with the DNA sequence containing the REF/ALT alleles, larger values are more significant. $E_{Diff}$ ($P_{REF}$ - $P_{ALT}$): Significance for allelic difference in p-values, calculated based on observed distribution of p-value differences in each of the 17 histone mark – cell type combinations. $E_{Diff}$ ($Sc_{REF}$ - $Sc_{ALT}$): Significance for allelic difference in scores, calculated based on observed distribution of score differences in each of the 17 histone mark – cell type combinations.

**Supplememntary Table S7: Deepbind transcription factor binding motifs that are affected by T1D SNPs that also affect thymocyte motifs.**

| | | SNP | motif allele | ID | $Sc_{REF}$ | $Sc_{ALT}$ | Diff | Protein | Type | Species | Family | Experiment | Experiment.Details |
|---|---|---|---|---|---|---|---|---|---|---|---|---|---|
| AAT1D | *PTPRK/ THEMIS* | rs138300818 | ALT | D00072.001 | 30,90 | 0,02 | 30,87 | Rfx7 | TF | *Mus musculus* | RFX | PBM | [DREAM5ID=TF_53, Array=ME] |
| AAT1D | *PTPRK/ THEMIS* | rs138300818 | ALT | D00619.003 | 5,13 | -1,96 | 7,09 | RFX5 | TF | *Homo sapiens* | RFX | SELEX | [CloneType=DBD, Primer=TGGAGC30NGAT, Cycle=4, Batch=AI] |
| AAT1D | *PTPRK/ THEMIS* | rs138300818 | ALT | D00616.002 | 4,07 | -1,44 | 5,51 | RFX3 | TF | *Homo sapiens* | RFX | SELEX | [CloneType=DBD, Primer=TGGCTT20NGA, Cycle=3, Batch=AC] |
| AAT1D | *PTPRK/ THEMIS* | rs138300818 | ALT | D00069.001 | 2,04 | 4,28 | 2,24 | Nr5a2 | TF | *Mus musculus* | Nuclear receptor | PBM | [DREAM5ID=TF_50, Array=ME] |
| AAT1D | *PTPRK/ THEMIS* | rs138300818 | ALT | D00579.002 | 0,98 | 3,07 | 2,09 | PHOX2A | TF | *Homo sapiens* | Homeo-domain | SELEX | [CloneType=DBD, Primer=TGACTC20NGA, Cycle=3, Batch=Y] |
| T1D | *AFF3* | rs66733041 | REF | D00054.001 | 3,30 | 0,19 | 3,11 | Atf3 | TF | *Mus musculus* | bZIP | PBM | [DREAM5ID=TF_35, Array=ME] |
| T1D | *AFF3* | rs66733041 | REF | D00410.003 | 3,24 | 0,95 | 2,29 | GATA3 | TF | *Homo sapiens* | GATA | SELEX | [CloneType=DBD, Primer=TGTCGT20NGA, Cycle=4, Batch=AC] |
| T1D | 7p15.2 | rs142852921 | REF | D00794.047 | 4,57 | 2,23 | 2,34 | POLR2A | TF | *Homo sapiens* | | ChIP-seq | [CellLine=MCF-7, Antibody=Pol2, Lab=UT-A] |

motif allele: allele that matches a thymocyte motif. ID: Deepbind transcription factor ID. Sc_REF/ Sc_ALT: Deepbind score for REF/ALT alleles; Diff: difference in REF/ALT scores. Protein: motif binding protein. Type: type of the protein, either transcription factor (TF) or RNA-bindig protein (RBP). Family: TF/RBP superfamily.

**Supplementary Table S8: Significant eQTL associations (FDP<0.05) in whole blood from the eQTLgen data base for SNPs in high LD (D'=1 and r2>0.89 in 1000Genomes GBP population) with rs138300818.** All SNPs belong to the 96% posterior probability credible set for association with age at diabetes diagnosis.

| Pvalue | SNP | SNPPos | Zscore | EA | NEA | Gene | NrCohorts | NrSamples | FDR |
|---|---|---|---|---|---|---|---|---|---|
| 9.45E-40 | rs7738609 | 128295502 | -13.1942 | T | C | *THEMIS* | 35 | 26482 | 0 |
| 2.83E-38 | rs9482850 | 128293506 | -12.9358 | T | C | *THEMIS* | 35 | 26494 | 0 |
| 2.95E-38 | rs72975916 | 128294055 | -12.9323 | T | C | *THEMIS* | 35 | 26493 | 0 |
| 3.47E-38 | rs9482851 | 128293634 | -12.9199 | T | C | *THEMIS* | 35 | 26493 | 0 |
| 3.66E-38 | rs72975913 | 128293932 | -12.9158 | A | C | *THEMIS* | 35 | 26493 | 0 |
| 4.32E-38 | rs72973800 | 128287158 | -12.9031 | T | C | *THEMIS* | 35 | 26494 | 0 |
| 5.26E-38 | rs11753289 | 128291681 | -12.8879 | G | T | *THEMIS* | 35 | 26488 | 0 |
| 6.73E-38 | rs9482849 | 128288536 | -12.8689 | C | T | *THEMIS* | 35 | 26486 | 0 |
| 8.04E-38 | rs72973797 | 128286386 | -12.855 | A | G | *THEMIS* | 35 | 26493 | 0 |
| 8.80E-38 | rs113297984 | 128286301 | -12.8481 | A | G | *THEMIS* | 35 | 26493 | 0 |
| 1.02E-37 | rs12111314 | 128289214 | -12.8366 | T | C | *THEMIS* | 35 | 26494 | 0 |
| 1.22E-37 | rs761332 | 128287848 | -12.8228 | A | G | *THEMIS* | 35 | 26494 | 0 |
| 2.52E-37 | rs9491893 | 128280931 | -12.7666 | A | G | *THEMIS* | 35 | 26494 | 0 |
| 3.44E-37 | rs9482848 | 128280375 | -12.7422 | T | A | *THEMIS* | 35 | 26494 | 0 |
| 4.76E-37 | rs6939352 | 128266250 | -12.717 | G | T | *THEMIS* | 35 | 26494 | 0 |
| 5.15E-37 | rs9491892 | 128280358 | -12.7107 | G | T | *THEMIS* | 35 | 26494 | 0 |
| 5.41E-37 | rs9491890 | 128270123 | -12.707 | A | G | *THEMIS* | 35 | 26494 | 0 |
| 7.59E-37 | rs4510698 | 128297611 | -12.6802 | T | C | *THEMIS* | 34 | 26156 | 0 |
| 7.85E-37 | rs9491891 | 128277151 | -12.6777 | G | A | *THEMIS* | 35 | 26493 | 0 |
| 1.16E-35 | rs118097399 | 128278233 | -12.4647 | T | C | *THEMIS* | 34 | 26408 | 0 |
| 5.57E-35 | rs3901020 | 128297604 | -12.3391 | G | C | *THEMIS* | 32 | 25563 | 0 |
| 1.94E-34 | rs9491889 | 128270067 | -12.2383 | T | C | *THEMIS* | 35 | 26494 | 0 |
| 1.14E-10 | rs9482849 | 128288536 | -6.4474 | C | T | *ECHDC1* | 34 | 30963 | 0 |
| 1.27E-10 | rs12111314 | 128289214 | -6.4308 | T | C | *ECHDC1* | 34 | 30971 | 0 |
| 1.30E-10 | rs72975913 | 128293932 | -6.427 | A | C | *ECHDC1* | 34 | 30970 | 0 |
| 1.36E-10 | rs11753289 | 128291681 | -6.4201 | G | T | *ECHDC1* | 34 | 30965 | 0 |
| 1.53E-10 | rs761332 | 128287848 | -6.4026 | A | G | *ECHDC1* | 34 | 30971 | 0 |
| 1.59E-10 | rs72973797 | 128286386 | -6.396 | A | G | *ECHDC1* | 34 | 30970 | 0 |
| 1.66E-10 | rs9482850 | 128293506 | -6.3902 | T | C | *ECHDC1* | 34 | 30971 | 0 |
| 1.68E-10 | rs9482851 | 128293634 | -6.388 | T | C | *ECHDC1* | 34 | 30970 | 0 |
| 1.70E-10 | rs113297984 | 128286301 | -6.3866 | A | G | *ECHDC1* | 34 | 30970 | 0 |
| 1.73E-10 | rs72973800 | 128287158 | -6.3834 | T | C | *ECHDC1* | 34 | 30971 | 0 |
| 2.10E-10 | rs72975916 | 128294055 | -6.3536 | T | C | *ECHDC1* | 34 | 30970 | 0 |
| 2.66E-10 | rs7738609 | 128295502 | -6.3171 | T | C | *ECHDC1* | 34 | 30959 | 0 |
| 2.69E-10 | rs118097399 | 128278233 | -6.3155 | T | C | *ECHDC1* | 33 | 30885 | 0 |
| 2.71E-10 | rs9491893 | 128280931 | -6.3146 | A | G | *ECHDC1* | 34 | 30971 | 0 |
| 2.89E-10 | rs9491892 | 128280358 | -6.3045 | G | T | *ECHDC1* | 34 | 30971 | 0 |
| 3.22E-10 | rs9482848 | 128280375 | -6.2879 | T | A | *ECHDC1* | 34 | 30971 | 0 |
| 3.69E-10 | rs9491889 | 128270067 | -6.2664 | T | C | *ECHDC1* | 34 | 30971 | 0 |
| 3.81E-10 | rs9491890 | 128270123 | -6.2617 | A | G | *ECHDC1* | 34 | 30971 | 0 |
| 3.81E-10 | rs9491891 | 128277151 | -6.2617 | G | A | *ECHDC1* | 34 | 30970 | 0 |
| 4.15E-10 | rs6939352 | 128266250 | -6.2481 | G | T | *ECHDC1* | 34 | 30971 | 0 |
| 7.67E-09 | rs3901020 | 128297604 | -5.7756 | G | C | *ECHDC1* | 31 | 30040 | 4.51E-05 |
| 1.30E-08 | rs4510698 | 128297611 | -5.6863 | T | C | *ECHDC1* | 33 | 30633 | 7.06E-05 |

EA: Effect allele, the allele in LD with the minor G insertion allele of rs138300818. NEA: non-effect allele. SNPpos: position on chromosome 6. NrCohorts/NrSamples: number of contributing cohorts/samples in eQTLgen data base . (www.eqtlgen.org)

**Supplementary Table S9: Thymocyte motifs overlapping type 1 diabetes credible set SNPs with marked difference (two orders of magnitude in p-value) in motif-sequence similarity between the reference and alternative alleles.** Significant motif overlap was considered as $P_{REF}$ or $P_{ALT} < 0.01/253$ thymocyte motifs/2467 autoimmune disease SNPs = $1.60\times10^{-8}$**.** Significant allelic difference was calculated based on observed distribution of p-value differences, separately for each of the 17 histone mark – cell type combinations and defined as $E_{Diff}$ ($P_{REF}$ - $P_{ALT}$) < 0.01/17 = $5.88\times10^{-4}$. Significant values are indicated with bold.

| SNP | REF | ALT | LOCUS | Matched Seq (REF/ALT) | Str | $P_{REF}$ | $P_{ALT}$ | $Sc_{REF}$ | $Sc_{ALT}$ | $E_{Diff}$ ($P_{REF}$ - $P_{ALT}$) | $E_{Diff}$ ($Sc_{REF}$ - $Sc_{ALT}$) | hmark | ctype | Count |
|---|---|---|---|---|---|---|---|---|---|---|---|---|---|---|
| rs142852921 | **G** | | 7p15.2 | GTAGTCCCAGCTACTTAGGAGGCTGAGG | + | ***4,49E-15*** | 5,04E-05 | 43,5 | -11,6 | ***5,4E-50*** | ***3,6E-07*** | state10 | CD3p_CD4p_CD8p | 12 |
| rs66733041 | **CTATGATGATAC** | | *AFF3* | AGTGCGGTGGTATCATCATAGCTCTCTGCA | - | ***1,35E-11*** | 6,72E-03 | 31,3 | -41,2 | ***3,0E-32*** | ***3,8E-04*** | H3K4me3 | CD4_ab | 5 |
| rs71024750 | | **GTGTGT** | *CENPW* | GTGTGTGTGTGTGTGTGTGTGTGTGTGTGT | + | ***8,43E-12*** | ***1,69E-18*** | 33,4 | 45,7 | ***2,7E-23*** | 4,5E-01 | state9 | CD3p_CD4p_CD8p | 4 |
| rs368755101 | G | | *AFF3* | TTTTTTTTTTTTTTTTTTTTT | - | 1,11E-07 | ***4,44E-13*** | 20,0 | 30,2 | ***4,3E-16*** | 4,4E-01 | H3K27ac | CD3p_CD4p_CD8p | 6 |
| rs588447 | **A** | C | *PTPN2* | TCTGTTGCCCAGGCTGGAGTGCAAT | + | ***2,76E-15*** | ***2,72E-10*** | 39,1 | 28,5 | ***3,6E-15*** | 3,3E-01 | state12 | CD3p_CD4p_CD8p | 8 |
| rs145917030 | | **GTTGTG** | *AFF3* | CTAAACACACACACACACACACACA | - | 3,06E-06 | ***4,90E-12*** | 5,0 | 34,3 | ***1,2E-14*** | 6,0E-02 | state12 | CD3n_CD4p_CD8p | 10 |
| rs79092647 | **AAA** | | *RBM17 IL2RA* | GGGCAAAAGTCTGTCTCAAAAAAAAAAAA | + | ***9,47E-14*** | 1,92E-08 | 38,6 | 20,8 | ***1,2E-13*** | 3,2E-01 | state11 | CD3p_CD4p_CD8p | 1 |
| rs7193670 | **T** | A | *DEXI CLEC16A* | GGGATTACAGGCATGAGCCACCGCGCCCG | - | ***1,57E-17*** | ***2,59E-13*** | 48,3 | 38,6 | ***6,8E-13*** | 4,6E-01 | state7 | CD3p_CD4p_CD8p | 9 |
| rs34361002 | | **AA** | *DEXI CLEC16A* | TAAAAAAAAAAAAAA | + | 2,48E-04 | ***7,15E-09*** | 2,1 | 23,8 | ***8,1E-13*** | 4,8E-02 | state12 | CD3p_CD4p_CD8p | 6 |
| rs9401891 | G | **T** | *CENPW* | TATTTTTCTTTTTTTTTTTTT | + | 3,88E-07 | ***1,11E-11*** | 18,4 | 27,4 | ***1,8E-11*** | 4,1E-01 | state10 | CD3p_CD4p_CD8p | 4 |
| rs67878610 | **TTT** | | *PTPN2* | ATTTATATTAAAATAAAAAAT | - | ***7,38E-11*** | 1,87E-06 | 26,4 | 16,0 | ***7,4E-11*** | 3,3E-01 | state10 | CD3p_CD4p_CD8p | 2 |
| rs3862471 | G | **T** | *DEXI CLEC16A* | AAACTCTGTCTCTACTAAAAATACAAAAAT | + | ***2,39E-11*** | ***1,80E-15*** | 31,0 | 39,7 | ***7,9E-11*** | 4,3E-01 | state12 | CD3p_CD4p_CD8p | 4 |
| rs571689 | **C** | T | *FUT2* | CCTGTAATCCCAGCTACTCGGGAGGCTGAG | + | ***1,42E-17*** | ***1,65E-13*** | 49,2 | 38,9 | ***1,5E-10*** | 3,4E-01 | state12 | CD3p_CD4p_CD8p | 4 |
| rs2309755 | G | **A** | *AFF3* | GGGATTACAGGCATGAGCCACCGCACCCA | + | ***1,58E-12*** | ***3,09E-16*** | 35,6 | 46,5 | ***2,1E-10*** | 3,9E-01 | state7 | CD3p_CD4p_CD8p | 6 |
| rs10175599 | C | **T** | *AFF3* | AGACTCTGTCTCAAAAAAAAAAAAAAAAG | - | ***4,75E-10*** | ***2,54E-14*** | 27,6 | 38,4 | ***2,6E-10*** | 3,2E-01 | state10 | CD3p_CD4p_CD8p | 11 |
| rs61634868 | **G** | C | *16q23.1* | GGGATTACAGGCATGCGCCACCACACCCA | - | ***3,98E-16*** | ***1,81E-12*** | 46,3 | 35,4 | ***4,7E-10*** | 4,1E-01 | state7 | CD3p_CD4p_CD8p | 5 |
| rs28665408 | **A** | C | *CD226* | TTATTTTCAATTTTTTTTTTTTTTTTTTTGT | - | ***2,93E-13*** | ***1,09E-08*** | 33,7 | 23,1 | ***5,3E-10*** | 6,0E-01 | H3K4me3 | CD4_ab | 9 |
| rs57071364 | **AATAAA** | | *14q24.1* | CTAATTATCTTTATTTTTATTTTTGTAGA | - | ***1,93E-09*** | 8,72E-05 | 25,3 | 6,5 | ***6,0E-10*** | 1,9E-01 | state12 | CD4_ab | 7 |
| rs6729966 | **A** | C | *AFF3* | AGCCGGGCGCGGTGGCTCACG | + | ***5,51E-12*** | 3,94E-08 | 30,7 | 21,2 | ***1,3E-09*** | 3,8E-01 | state12 | CD3p_CD4p_CD8p | 5 |
| rs147034755 | A | **C** | *FAM98B; SPRED1; RASGRP1* | GGGATTACAGGCATGTGCCACCATGCCCG | - | ***2,64E-12*** | ***8,47E-16*** | 34,7 | 45,6 | ***2,2E-09*** | 3,9E-01 | state7 | CD3p_CD4p_CD8p | 5 |
| rs1790962 | **T** | G | *CD226* | TCTCGCTCTGTCGCCCAGGCTGGAGTGC | + | ***7,86E-16*** | ***2,02E-11*** | 43,2 | 32,1 | ***3,3E-09*** | 5,8E-01 | H3K4me3 | CD3n_CD4p_CD8p | 10 |
| rs1295793 | **C** | G | *14q24.1* | CCTGTAATCCCAGCATTTTGGGAGGCTGAG | - | ***1,05E-16*** | ***5,49E-13*** | 47,3 | 37,2 | ***4,7E-09*** | 3,5E-01 | state12 | CD3p_CD4p_CD8p | 3 |
| rs4851259 | A | **C** | *AFF3* | CTGTAATCCCAGCACTTTGGGAGGCTGAGG | + | ***5,72E-14*** | ***2,02E-18*** | 40,7 | 50,7 | ***6,1E-09*** | 6,4E-01 | H3K27ac | CD3n_CD4p_CD8p | 11 |
| rs1688264 | **T** | G | *FUT2* | AAAAAAAAAAAAAAT | + | ***1,35E-10*** | 2,97E-06 | 25,2 | 15,1 | ***6,6E-09*** | 5,0E-01 | state12 | CD3n_CD4p_CD8p | 6 |
| rs11900482 | G | **C** | *AFF3* | ACACACACATGCACACGCACACACACTCA | + | ***1,09E-09*** | ***2,59E-13*** | 26,4 | 36,3 | ***1,1E-08*** | 3,7E-01 | state12 | CD3p_CD4p_CD8p | 3 |
| rs147955926 | **TT** | | *14q32.2* | AAAAAAAAAAAAAAA | - | ***3,49E-10*** | 1,31E-06 | 26,9 | 16,0 | ***1,8E-08*** | 3,1E-01 | state12 | CD3p_CD4p_CD8p | 14 |
| rs1989265 | T | **G** | *COBL* | TGAACCTGGGAGGCGAAGGTTGCAGTGAGC | + | ***9,60E-13*** | ***2,98E-16*** | 36,6 | 46,2 | ***3,2E-08*** | 3,8E-01 | state12 | CD3p_CD4p_CD8p | 3 |
| rs3823931 | **T** | C | *7p15.2* | ATTTAAAGTATGTAAAGAAGA | - | ***2,18E-10*** | 2,65E-06 | 25,7 | 15,4 | ***4,8E-08*** | 5,0E-01 | state12 | CD3n_CD4p_CD8p | 1 |

| SNP | REF | ALT | LOCUS | Matched Seq (REF/ALT) | Str | $P_{REF}$ | $P_{ALT}$ | $Sc_{REF}$ | $Sc_{ALT}$ | $E_{Diff}$ ($P_{REF}$ - $P_{ALT}$) | $E_{Diff}$ ($Sc_{REF}$ - $Sc_{ALT}$) | hmark | ctype | Count |
|---|---|---|---|---|---|---|---|---|---|---|---|---|---|---|
| rs12957037 | **T** | G | *PTPN2* | TTTTTGTATTTTTTGTAGAGACAGAGTTT | + | ***8,19E-17*** | ***1,16E-12*** | 47,3 | 36,2 | ***4,8E-08*** | 4,6E-01 | H3K27me3 | CD3n_CD4p_CD8p | 2 |
| rs7785832 | **T** | C | *7p15.2* | ATTTTTTTAAAAATTATTAGTTAAATTTT | + | ***7,55E-10*** | 3,45E-06 | 25,6 | 14,5 | ***6,2E-08*** | 3,1E-01 | state10 | CD3p_CD4p_CD8p | 1 |
| rs692854 | C | **A** | *FUT2* | TGAACCAGGGAGGCAGAGGTTGCAGTGAGC | + | ***2,36E-12*** | ***1,03E-15*** | 35,2 | 45,0 | ***1,2E-07*** | 3,7E-01 | state12 | CD3p_CD4p_CD8p | 4 |
| rs73067437 | **T** | C | *7p15.2* | TTTGTTTGTTTTTTT | + | ***1,67E-09*** | 6,06E-06 | 24,2 | 13,9 | ***1,3E-07*** | 5,1E-01 | state9 | CD3p_CD4p_CD8p | 4 |
| rs1985869 | **C** | G | *DEXI CLEC16A* | CCTGGCCTCAAGTGATCCTCC | + | ***1,23E-12*** | ***2,51E-09*** | 34,3 | 25,3 | ***1,9E-07*** | 4,1E-01 | state12 | CD3p_CD4p_CD8p | 4 |
| rs111279202 | **T** | | *DEXI CLEC16A* | TTTTTTTTCTTTTTTTTTTTT | + | ***1,81E-14*** | ***1,67E-10*** | 33,6 | 28,2 | ***1,9E-07*** | 7,1E-01 | H3K27me3 | CD3n_CD4p_CD8p | 4 |
| rs12712070 | **C** | A | *AFF3* | TTGTTGCCCAGGCTGGAGTGGAATGG | - | ***3,13E-13*** | ***9,52E-10*** | 36,9 | 26,4 | ***2,6E-07*** | 3,3E-01 | state10 | CD3p_CD4p_CD8p | 3 |
| rs2452170 | **G** | A | *FUT2* | CCTGTAGTTCCAGCTACTCGGGAGGCTGAG | - | ***2,58E-15*** | ***4,76E-12*** | 44,2 | 33,9 | ***2,7E-07*** | 3,4E-01 | state12 | CD3p_CD4p_CD8p | 2 |
| rs507711 | **C** | T | *FUT2* | CTGACCAACATGGTGAAACTCCATTTCCAC | - | ***7,07E-14*** | ***4,96E-10*** | 36,2 | 27,5 | ***2,8E-07*** | 5,6E-01 | state12 | CD3n_CD4p_CD8p | 1 |
| rs139377078 | AGAG | | *7p15.2* | ATATTTTTCCATTTGTATGTGTTCTTATTT | - | 1,53E-05 | ***2,04E-09*** | 8,5 | 25,4 | ***2,8E-07*** | 2,5E-01 | state12 | CD4_ab | 1 |
| rs1988588 | **T** | A | *14q32.2* | TGAGACGGAGTCTCGCTCTGTTGCCCAGG | - | ***4,16E-16*** | ***1,18E-12*** | 44,7 | 36,3 | ***2,9E-07*** | 5,9E-01 | state9 | CD3p_CD4p_CD8p | 4 |
| rs507855 | A | **G** | *FUT2* | TCCCAGCTACTCGGGAGGCTGAGGCAGGAG | - | ***4,32E-16*** | ***4,42E-19*** | 46,0 | 57,1 | ***3,0E-07*** | 3,8E-01 | state7 | CD3p_CD4p_CD8p | 1 |
| rs1702877 | **C** | T | *IKZF4 DGKA ERBB3* | TCCCAGCTACTCGGGAGGCTGAGGCAGGAG | - | ***4,42E-19*** | ***4,32E-16*** | 57,1 | 46,0 | ***3,6E-07*** | 4,0E-01 | state7 | CD3p_CD4p_CD8p | 2 |
| rs6715254 | **C** | A | *AFF3* | CTGTAATCCCAGCACTTTGGGAGGCCGAGG | + | ***2,74E-17*** | ***2,06E-13*** | 48,9 | 38,8 | ***3,7E-07*** | 6,2E-01 | H3K27ac | CD3n_CD4p_CD8p | 6 |
| rs13207431 | **C** | T | *CENPW* | TCACGCCATTCTCCTGCCTCAGCCTCCCA | + | ***1,25E-15*** | ***1,14E-12*** | 46,1 | 35,0 | ***4,8E-07*** | 4,0E-01 | state7 | CD3p_CD4p_CD8p | 2 |
| rs60455438 | C | **T** | *FAM98B; SPRED1; RASGRP1* | TCTGTCACCCAGGCTGGAGTGCAGT | + | ***1,46E-09*** | ***9,37E-13*** | 25,6 | 36,1 | ***4,8E-07*** | 3,4E-01 | state12 | CD3p_CD4p_CD8p | 5 |
| rs34813703 | GTG | | *IKZF4 DGKA ERBB3* | TGACACGCCTGTAATCCCAGTACTTC | - | 1,55E-05 | ***7,08E-09*** | -0,8 | 21,4 | ***4,9E-07*** | 9,0E-02 | H3K27ac | CD3p_CD4p_CD8p | 1 |
| rs113003633 | C | **T** | *7p15.2* | AGTGCAGTGGCATGATCTCGGCTCACTGCA | - | ***2,40E-13*** | ***5,21E-17*** | 38,3 | 49,0 | ***6,0E-07*** | 6,0E-01 | H3K4me3 | CD4_ab | 5 |
| rs1262550 | **A** | C | *CENPW* | AAACAAAAAAAAACA | + | ***3,19E-09*** | 7,20E-06 | 23,2 | 12,3 | ***7,1E-07*** | 3,1E-01 | state10 | CD3p_CD4p_CD8p | 9 |
| rs3781196 | **G** | T | *RNLS* | TCTGTGCCTTGGTTTCCTCACTTGTTAAA | + | ***1,63E-10*** | 2,02E-07 | 28,7 | 18,2 | ***1,1E-06*** | 3,3E-01 | state12 | CD3p_CD4p_CD8p | 1 |
| rs4897180 | **A** | T | *CENPW* | ATCTTGGCTCACTGCAACCTCCGCCTCC | - | ***9,23E-17*** | ***5,62E-13*** | 46,7 | 37,6 | ***1,6E-06*** | 4,9E-01 | H3K27ac | CD4_ab | 4 |
| rs12923098 | **T** | C | *DEXI CLEC16A* | GCCTGGGCAACAGAGCAAGAC | - | ***1,75E-12*** | ***7,63E-09*** | 33,9 | 23,5 | ***2,0E-06*** | 4,9E-01 | H3K4me3 | CD8_ab | 2 |
| rs59254259 | **T** | G | *7p15.2* | ACTGCACTCCAGCCTGGGCGA | + | ***3,00E-13*** | ***6,98E-10*** | 36,9 | 27,2 | ***2,5E-06*** | 5,9E-01 | state11 | CD3p_CD4p_CD8p | 7 |
| rs1985872 | G | **C** | *DEXI CLEC16A* | GCTCACGCCTGTAATTCCAGCACTTT | - | ***5,61E-10*** | ***4,24E-13*** | 26,6 | 37,4 | ***2,6E-06*** | 4,1E-01 | H3K27ac | CD3p_CD4p_CD8p | 5 |
| rs11899489 | **A** | C | *AFF3* | CCTGGGCTCAAGTGATCCTCC | - | ***1,58E-12*** | ***1,51E-09*** | 34,1 | 26,2 | ***2,7E-06*** | 4,7E-01 | state12 | CD3p_CD4p_CD8p | 3 |
| rs4851260 | C | **A** | *AFF3* | GGGATTACAGGCACGTGCCACCATGCCCA | - | ***4,44E-12*** | ***8,72E-15*** | 33,7 | 43,2 | ***3,5E-06*** | 4,5E-01 | state7 | CD3p_CD4p_CD8p | 6 |
| rs507766 | **T** | C | *FUT2* | TAAACATATAAAAAT | - | ***3,14E-09*** | 9,18E-06 | 23,7 | 12,6 | ***3,6E-06*** | 4,6E-01 | state12 | CD3n_CD4p_CD8p | 2 |
| rs7192287 | A | **T** | *DEXI CLEC16A* | GGGATTACAGGCGCCTGCCACCATGCCTG | + | ***1,62E-11*** | ***3,23E-14*** | 31,2 | 41,6 | ***3,6E-06*** | 4,1E-01 | state7 | CD3p_CD4p_CD8p | 4 |
| rs10227673 | G | **A** | *7p15.2* | ATTTTTATTATTTTT | - | 2,12E-06 | ***4,80E-10*** | 15,4 | 25,6 | ***4,3E-06*** | 4,6E-01 | H3K27ac | CD4_ab | 3 |
| rs2548459 | **T** | C | *FUT2* | AACTCCTGACCTCAGGTGATC | - | ***2,16E-13*** | ***5,59E-10*** | 37,2 | 27,5 | ***4,7E-06*** | 6,3E-01 | H3K4me3 | CD3n_CD4p_CD8p | 6 |
| rs28894750 | A | **T** | *FUT2* | AAAAAAAAATTAGCCAGGTGTG | - | 3,94E-08 | ***2,26E-11*** | 20,6 | 31,4 | ***5,8E-06*** | 5,6E-01 | state11 | CD3p_CD4p_CD8p | 1 |
| rs506897 | **G** | C | *FUT2* | CCTGTAATCCCAGCACTTTGGGAGGCCGA | - | ***2,34E-17*** | ***5,58E-14*** | 50,5 | 40,6 | ***5,9E-06*** | 6,2E-01 | H3K4me3 | CD3n_CD4p_CD8p | 1 |
| rs6564237 | C | **G** | *16q23.1* | AGGAAGATCGCTTGAGCCCAG | - | 3,52E-08 | ***3,14E-11*** | 21,3 | 30,6 | ***6,3E-06*** | 3,9E-01 | state10 | CD3p_CD4p_CD8p | 2 |

| SNP | REF | ALT | LOCUS | Matched Seq (REF/ALT) | Str | $P_{REF}$ | $P_{ALT}$ | $Sc_{REF}$ | $Sc_{ALT}$ | $E_{Diff}$ ($P_{REF}$ - $P_{ALT}$) | $E_{Diff}$ ($Sc_{REF}$ - $Sc_{ALT}$) | hmark | ctype | Count |
|---|---|---|---|---|---|---|---|---|---|---|---|---|---|---|
| rs28816386 | A | **G** | *7p15.2* | GGCCTCCCAAAGTGCTGGGATTACAGGCG | - | ***3,18E-14*** | ***1,33E-17*** | 41,7 | 50,1 | ***7,7E-06*** | 5,7E-01 | state12 | CD4_ab | 2 |
| rs3030572 | | **A** | *DEXI CLEC16A* | AGCTAATTTTTTTTTTAATTAAAATTATCT | + | 5,92E-08 | ***2,56E-11*** | 20,0 | 31,1 | ***8,4E-06*** | 4,5E-01 | state12 | CD4_ab | 2 |
| rs4930045 | **T** | A | *INS* | CCATCCTGGCTAACACAGTGAAACCCCGTC | - | ***9,75E-13*** | ***1,85E-09*** | 35,8 | 25,1 | ***8,5E-06*** | 6,0E-01 | H3K4me3 | CD4_ab | 2 |
| rs75169667 | **A** | G | *CENPW* | AAGAAGGAAAGAGAG | + | ***1,33E-08*** | 2,76E-05 | 21,6 | 10,4 | ***9,2E-06*** | 4,6E-01 | state12 | CD3n_CD4p_CD8p | 1 |
| rs9388496 | **A** | G | *CENPW* | TTATTTAAAAATTTT | - | ***3,67E-09*** | 9,15E-06 | 23,0 | 12,9 | ***9,2E-06*** | 5,0E-01 | H3K4me3 | CD8_ab | 1 |
| rs3850234 | **T** | A | *CENPW* | GTAATCCTAGCAATTTGGGAGGATGAGG | - | ***3,83E-14*** | ***3,77E-11*** | 41,1 | 30,1 | ***9,6E-06*** | 3,1E-01 | state10 | CD3p_CD4p_CD8p | 1 |
| rs71136618 | | **T** | *DEXI CLEC16A* | TTTTTTTTTTCTTTTTTTTTT | + | ***7,80E-09*** | ***3,37E-12*** | 23,8 | 31,2 | ***1,0E-05*** | 6,4E-01 | H3K27me3 | CD3n_CD4p_CD8p | 4 |
| rs1788100 | **A** | G | *CD226* | AAAAAAAAAAAAATAT | + | ***1,90E-09*** | 3,61E-06 | 24,0 | 14,7 | ***1,2E-05*** | 5,3E-01 | state12 | CD3n_CD4p_CD8p | 4 |
| rs2214494 | G | **T** | *COBL* | ATTTTTCCTTTTCACTTAATATTTTTAAAT | + | 6,69E-07 | ***4,25E-10*** | 16,7 | 27,1 | ***1,3E-05*** | 6,1E-01 | H3K4me3 | CD4_ab | 1 |
| rs570794 | T | **C** | *FUT2* | TGAGATGGAGTCTTGCTGTGTCACCCAGG | - | ***3,70E-11*** | ***4,42E-14*** | 30,9 | 40,4 | ***1,4E-05*** | 5,6E-01 | state9 | CD3p_CD4p_CD8p | 3 |
| rs3861458 | G | **A** | *CENPW* | TTTAAATTTAAGTTTACATAATGGATTAT | - | 2,32E-06 | ***1,27E-09*** | 15,3 | 25,1 | ***1,4E-05*** | 5,4E-01 | state12 | CD3n_CD4p_CD8p | 1 |
| rs111869668 | **C** | G | *16q23.1* | CTGGAGTGCAGTGGTGCAATCTTGGCTCC | - | ***2,37E-14*** | ***1,75E-11*** | 41,5 | 31,8 | ***2,1E-05*** | 5,3E-01 | state9 | CD3p_CD4p_CD8p | 4 |
| rs8054218 | **A** | G | *16q23.1* | GCCTGGGTGACAGAGTGAGAC | + | ***1,43E-11*** | 2,39E-08 | 32,1 | 21,5 | ***2,6E-05*** | 4,8E-01 | H3K4me3 | CD8_ab | 2 |
| rs145665108 | TGAAAT | | *7p15.2* | AAATAAATGAAATTTTAAAAAAATACAAAA | + | 7,23E-08 | ***7,79E-11*** | 18,0 | 30,2 | ***3,0E-05*** | 4,5E-01 | H3K4me3 | CD3p_CD4p_CD8p | 4 |
| rs5815611 | **T** | | *DEXI CLEC16A* | TTTCTTTTTTTTTTT | + | ***7,12E-10*** | 8,41E-07 | 25,7 | 16,9 | ***3,8E-05*** | 6,6E-01 | H3K4me3 | CD3n_CD4p_CD8p | 1 |
| rs112166936 | A | **C** | *CENPW* | CCTCCCTTCCTCTCTCCCTCC | + | 9,09E-08 | ***7,71E-11*** | 20,4 | 28,5 | ***3,8E-05*** | 7,0E-01 | H3K4me3 | CD3n_CD4p_CD8p | 1 |
| rs28362844 | C | | *FUT2* | ATGTTTTCTTTTCTTTTTCTTTTCTTTTT | + | 6,06E-06 | ***1,06E-08*** | 13,4 | 22,9 | ***4,5E-05*** | 3,9E-01 | state10 | CD3p_CD4p_CD8p | 1 |
| rs1967332 | **C** | A | *7p15.2* | CCACTGCACCCCAGCCTGGGTGACAAAG | + | ***9,35E-15*** | ***4,60E-12*** | 43,2 | 33,3 | ***4,9E-05*** | 4,4E-01 | H3K27ac | CD3p_CD4p_CD8p | 2 |
| rs9952753 | A | **C** | *PTPN2* | GGGATTACAGGCGTAAACCACCCCGCCCA | + | ***7,74E-11*** | ***3,33E-13*** | 27,8 | 38,2 | ***4,9E-05*** | 4,1E-01 | state7 | CD3p_CD4p_CD8p | 1 |
| rs10758593 | G | **A** | *GLIS3* | AAAAAGATTTTTAAGAAAAACATAAATAA | + | 2,23E-06 | ***2,01E-09*** | 15,3 | 24,6 | ***5,1E-05*** | 5,6E-01 | state12 | CD3n_CD4p_CD8p | 1 |
| rs35536837 | | **T** | *CENPW* | AAAAAAAAAGGAAAAGAAAAATTATATCA | - | 6,89E-06 | ***9,17E-09*** | 12,6 | 23,2 | ***5,8E-05*** | 5,7E-01 | state11 | CD3p_CD4p_CD8p | 1 |
| rs8180649 | T | **C** | *CENPW* | GAACTCCTGACCTCAGGTGAT | + | ***1,11E-09*** | ***9,33E-13*** | 26,6 | 35,4 | ***6,0E-05*** | 6,8E-01 | H3K27ac | CD3n_CD4p_CD8p | 2 |
| rs12592898 | A | **G** | *CTSH* | AACTCCCGACCTCAGGTGATC | + | ***1,74E-09*** | ***1,83E-12*** | 25,6 | 34,9 | ***6,5E-05*** | 6,5E-01 | H3K4me3 | CD3n_CD4p_CD8p | 1 |
| rs139078271 | | **G** | *CENPW* | TATAATTTCTGTTCTTTTACATTTGCTGA | - | 5,19E-07 | ***5,28E-10*** | 17,1 | 26,9 | ***7,4E-05*** | 5,1E-01 | state12 | CD4_ab | 1 |
| rs112164743 | **A** | G | *7p15.2* | AGAACTTAAAGTATACTTAAAAAAAAAAA | + | ***5,15E-09*** | 6,73E-06 | 23,7 | 13,2 | ***7,9E-05*** | 4,3E-01 | H3K27ac | CD4_ab | 1 |
| rs67518044 | **TAT** | | *PTPN2* | AAACAAAACAAATAATACCACAATAAAAA | - | ***7,19E-10*** | 6,70E-07 | 26,9 | 15,0 | ***1,1E-04*** | 4,3E-01 | H3K4me3 | CD8_ab | 1 |
| rs9939397 | **A** | G | *DEXI CLEC16A* | ATATATTATAAATTTTTTTAAAGACAAGG | + | ***3,48E-09*** | 2,80E-06 | 24,5 | 14,0 | ***1,1E-04*** | 4,6E-01 | state12 | CD4_ab | 1 |
| rs1704773 | **A** | G | *FUT2* | ACTCTTAAGAATTTTTTTTTTTTTTTTTGA | - | ***9,82E-09*** | 2,78E-06 | 23,0 | 12,4 | ***1,1E-04*** | 3,3E-01 | state12 | CD3p_CD4p_CD8p | 2 |
| rs2847291 | **G** | A | *PTPN2* | GGAGGCGGAGGTTGC | + | ***1,84E-09*** | 1,66E-06 | 26,2 | 15,4 | ***1,1E-04*** | 4,8E-01 | H3K4me3 | CD8_ab | 1 |
| rs184458383 | **C** | T | *16q23.1* | TTAGATTCGGCTGGGTGTGGTGGCTCACGC | + | ***3,37E-11*** | 2,06E-08 | 31,5 | 20,7 | ***1,6E-04*** | 6,0E-01 | H3K4me3 | CD4_ab | 1 |
| rs377093688 | **C** | G | *FAM98B SPRED1 RASGRP1* | GCGATCTCGGCTCACTGCAA | + | ***1,30E-12*** | ***8,49E-10*** | 36,5 | 26,6 | ***1,6E-04*** | 6,2E-01 | H3K4me3 | CD3n_CD4p_CD8p | 1 |
| rs8056098 | G | **A** | *DEXI CLEC16A* | TGGCAATTAGTTTTTTTTTTTTTTTTTTTGA | - | 3,47E-07 | ***4,98E-10*** | 16,8 | 27,3 | ***1,7E-04*** | 4,8E-01 | state12 | CD4_ab | 1 |
| rs7184615 | G | **C** | *DEXI CLEC16A* | CCTGGGCTCAAGTGATCATCC | + | ***1,25E-08*** | ***5,32E-11*** | 22,0 | 30,9 | ***1,8E-04*** | 4,1E-01 | state12 | CD3p_CD4p_CD8p | 3 |

| SNP | REF | ALT | LOCUS | Matched Seq (REF/ALT) | Str | $P_{REF}$ | $P_{ALT}$ | $Sc_{REF}$ | $Sc_{ALT}$ | $E_{Diff}$ $(P_{REF} - P_{ALT})$ | $E_{Diff}$ $(Sc_{REF} - Sc_{ALT})$ | hmark | ctype | Count |
|---|---|---|---|---|---|---|---|---|---|---|---|---|---|---|
| rs141552689 | | **AA** | *7p15.2* | ATTTTTTTTTTAAATTAAAACAAGTTTT | - | 7,26E-07 | ***1,01E-09*** | 10,8 | 25,9 | ***1,9E-04*** | 3,2E-01 | H3K4me3 | CD8_ab | 1 |
| rs9926367 | **T** | C | *DEXI CLEC16A* | AAGTGTGCTTGTGTGGGTGTG | + | ***1,37E-08*** | 7,99E-06 | 22,8 | 12,0 | ***2,1E-04*** | 5,9E-01 | H3K4me3 | CD3n_CD4p_CD8p | 1 |
| rs75407602 | C | **T** | *TYK2* | TTTGATTTTTTTTTTTTTTTCCAGAGATGG | + | 2,18E-06 | ***3,43E-09*** | 14,5 | 24,6 | ***2,1E-04*** | 4,9E-01 | state12 | CD4_ab | 1 |
| rs34752850 | **A** | | *DEXI CLEC16A* | ATCTTCTTTTTTTTTAAGAAACAAGGTCT | - | ***4,10E-09*** | 2,55E-06 | 20,5 | 2,1 | ***2,4E-04*** | 2,2E-01 | H3K27me3 | CD3n_CD4p_CD8p | 1 |
| rs12416116 | C | **A** | *RNLS* | ATACCTCATTAGAACAACAAAAAAAAAAT | + | 9,55E-07 | ***3,25E-09*** | 13,8 | 24,7 | ***2,6E-04*** | 3,2E-01 | state10 | CD3p_CD4p_CD8p | 1 |
| rs5823237 | T | | *PTPN2* | ATCGTAGGTCACTGCAGACTCAAACTCC | - | 9,55E-07 | ***1,24E-09*** | 1,3 | 21,5 | ***2,8E-04*** | 1,4E-01 | H3K27ac | CD4_ab | 2 |
| rs2847292 | **G** | T | *PTPN2* | ATGGGGTTTCACCATGTTGGCCAGGCTGGT | + | ***6,79E-15*** | ***4,89E-12*** | 43,1 | 34,0 | ***2,9E-04*** | 4,9E-01 | H3K27ac | CD4_ab | 1 |
| rs28404962 | **T** | C | *7p15.2* | AAAAGTTGAATATTAAATCATAAAAATAA | + | ***1,13E-08*** | 7,50E-06 | 22,7 | 13,2 | ***3,5E-04*** | 4,7E-01 | H3K27ac | CD4_ab | 1 |
| rs12722552 | **C** | T | *RBM17 IL2RA* | TGGTCTCGAACTCCTGACCTCAGGTGATC | + | ***3,54E-18*** | ***1,65E-15*** | 54,0 | 46,1 | ***3,6E-04*** | 6,0E-01 | state12 | CD3n_CD4p_CD8p | 1 |
| rs2847273 | **A** | C | *PTPN2* | TTTTTGTATTTTTAGTAGAGATGGAGTTG | - | ***8,84E-13*** | ***4,46E-10*** | 36,6 | 25,5 | ***3,8E-04*** | 4,6E-01 | H3K27me3 | CD3n_CD4p_CD8p | 1 |
| rs443081 | **C** | T | *19q13.32* | TCCCAGCTACTTGGGAGGCTGAGGTAGGAG | - | ***1,84E-17*** | ***2,23E-15*** | 53,5 | 42,4 | ***4,0E-04*** | 4,0E-01 | state7 | CD3p_CD4p_CD8p | 1 |
| rs1120786 | **T** | G | *CENPW* | TTGTTTTATTGTAAATCGTTATTTTATTAT | + | ***1,68E-09*** | 6,65E-07 | 25,5 | 16,7 | ***4,2E-04*** | 6,7E-01 | H3K4me3 | CD4_ab | 1 |
| rs2292759 | A | **G** | *PTPN2* | CCCGCCCCGCCGCCGACTCCGCGCCGCGC | - | 1,58E-06 | ***9,26E-09*** | 14,5 | 23,4 | ***4,3E-04*** | 4,2E-01 | state12 | CD3p_CD4p_CD8p | 1 |
| rs194739 | **C** | T | *14q24.1* | TCCCAGCTACTCGGGAGGCTGAGGCAGGGG | - | ***3,64E-17*** | ***4,03E-15*** | 52,1 | 41,0 | ***5,2E-04*** | 4,0E-01 | state7 | CD3p_CD4p_CD8p | 2 |
| rs200395517 | | **T** | *19q13.32* | ATTTTTTTTTTCTTTTTCTTTTTATTGTTT | + | 1,71E-08 | ***4,86E-11*** | 22,4 | 29,3 | ***5,2E-04*** | 7,3E-01 | H3K4me3 | CD4_ab | 1 |
| rs11680485 | **A** | G | *AFF3* | TCTAAAAAATAAAATAAAATA | + | ***5,74E-10*** | 2,16E-07 | 27,7 | 16,6 | ***5,5E-04*** | 5,8E-01 | H3K4me3 | CD3n_CD4p_CD8p | 1 |

Count: number of histone modification/ state – cell type combinations where there was significant allelic difference ($p<5.88\times10^{-4}$), out of maximum 17.

**Supplementary Figure S1: overlap between the discovered thymocyte motifs and known transcription factor (TF) binding motifs. For each TF, only the thymocyte motif from the cell type and histone modification peak combination with the lowest q-value is shown.**

**A: ZNF263, H3K4me3: CD3- CD4+ CD8+ *q*=1.35×10$^{-7}$**

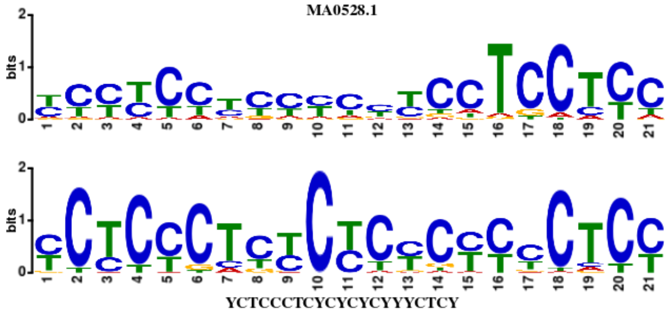

**B: SP2, H3K4me3: CD3- CD4+ CD8+ *q*=1.32×10$^{-6}$**

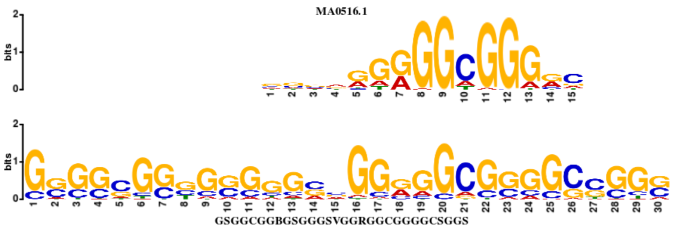

**C: SP1, H3K4me3: CD3- CD4+ CD8+ *q*=7.19×10$^{-5}$**

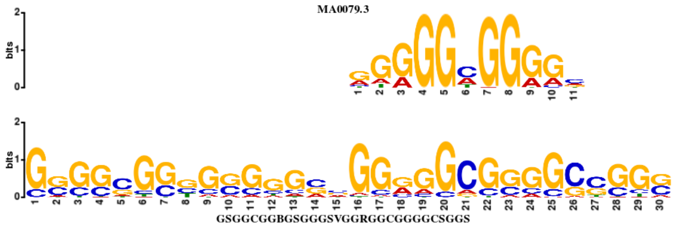

**D: Zfp281 (Znf281), H3K27ac: CD3+ CD4+ CD8+ *q*=3.75×10$^{-4}$**

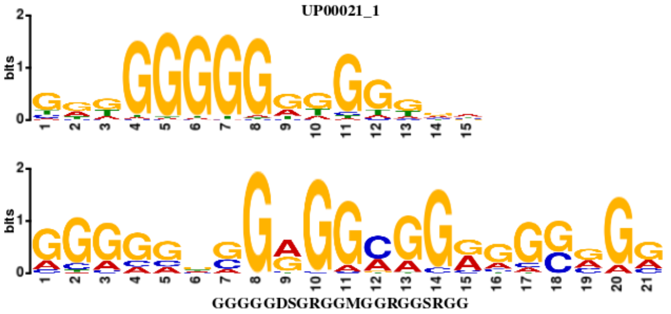

**E: SP4, H3K4me3: CD3- CD4+ CD8+ *q*=5.36×10$^{-4}$**

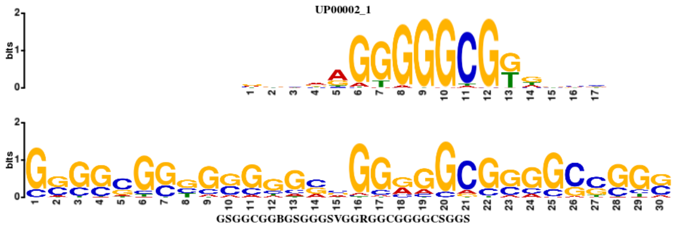

**F: KLF16, H3K4me3: CD8+ αß *q*=9.77×10$^{-4}$**

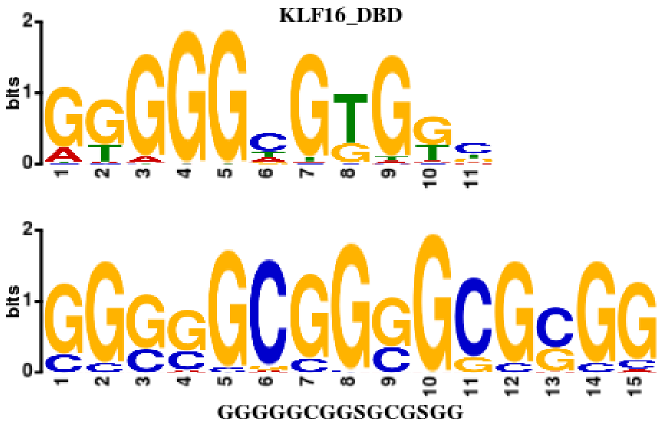

**G: KLF5, H3K4me3: CD8+ αß *q*=2.43×10$^{-3}$**

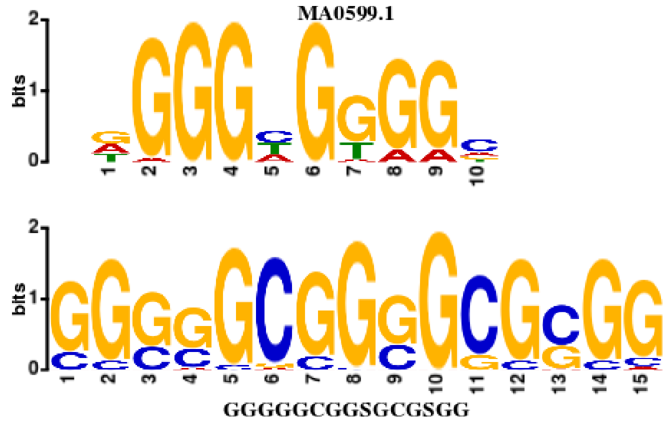

**H: SP3, H3K4me3: CD8+ αß *q*=3.04×10$^{-3}$**

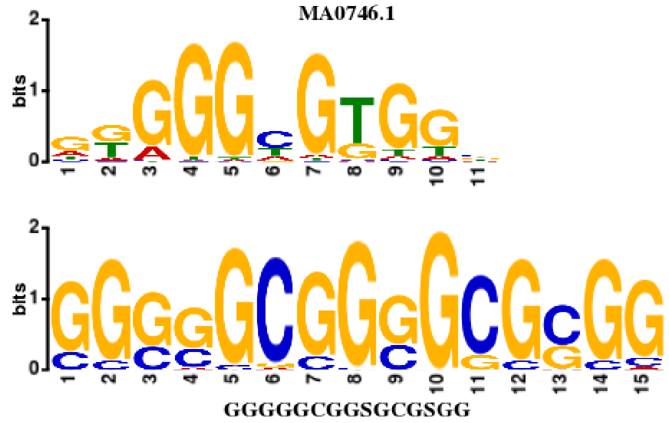

**I: RREB1, Stete12: CD3- CD4+ CD8+ $q$=3.85×10$^{-3}$**

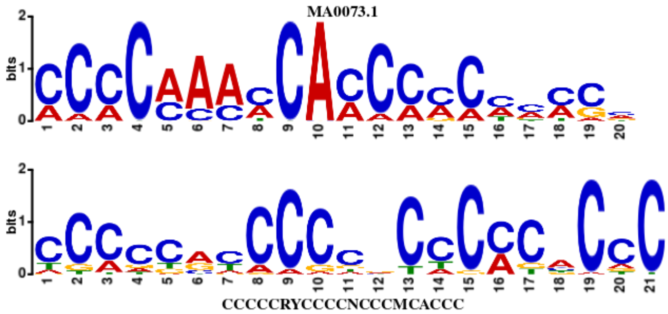

**J: Zfx, H3K4me3: CD8+ αß $q$=4.17×10$^{-3}$**

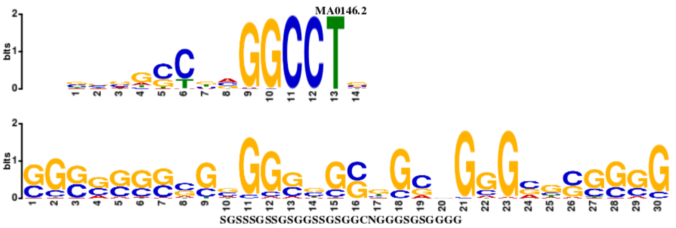

**K: Zfp740 (Znf740), H3K4me3: CD8+ αß $q$=9.11×10$^{-3}$**

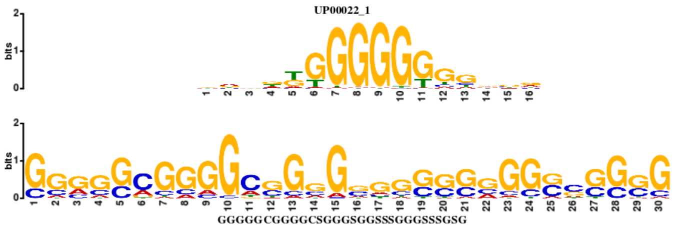

**Supplementary Figure S2: Predicted TF binding for all SNPs in credible set for age at diabetes diagnosis based on deepbind neural network predictions. A:** X-axis reports binding for REF allele, y-axis for the ALT allele. SNP – TF pairs with largest difference between alleles are annotated. **B:** Table of all SNP - TF pairs where allelic difference was larger than 5 standard deviations (≥6.43) of difference distribution at least with one TF binding; differences ≥6.43 are highlighted with bold.

**A**

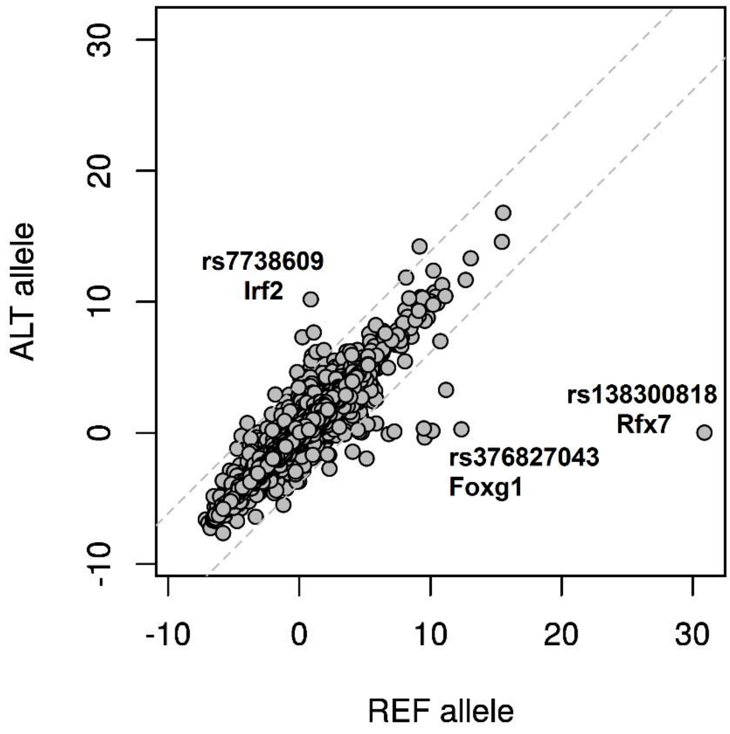

**B**

| TF ID | TF name | rs13204742 | | rs802747 | | rs376827043 | | rs1089652 | | rs7738609 | | rs138300818 | |
|---|---|---|---|---|---|---|---|---|---|---|---|---|---|
| | | REF G | ALT T | REF G | ALT A | REF GTTT | ALT - | REF C | ALT T | REF C | ALT T | REF - | ALT G |
| D00201.001 | CNOT4 | **0.238** | **7.301** | -0.167 | -0.191 | 0.026 | -0.080 | 4.868 | 4.868 | **4.751** | **2.693** | -0.101 | -0.143 |
| D00061.001 | Foxc2 | -0.286 | -0.366 | 0.071 | 0.060 | **9.498** | **0.340** | **1.874** | **6.322** | 1.255 | 1.143 | 0.184 | 0.178 |
| D00062.001 | Foxg1 | -0.459 | -0.140 | 0.328 | 0.449 | **12.364** | **0.273** | **1.100** | **7.664** | 0.495 | 0.476 | -0.654 | -0.526 |
| D00004.001 | Foxj2 | -0.457 | -0.439 | -0.283 | -0.286 | **9.560** | **-0.352** | -0.072 | 0.585 | 0.669 | 0.390 | -0.539 | -0.486 |
| D00005.001 | Foxo1 | -0.166 | -0.166 | 0.232 | 0.000 | **6.819** | **-0.091** | 0.828 | 1.424 | 0.191 | 0.172 | -0.161 | -0.162 |
| D00008.001 | Foxp1 | -0.083 | -0.083 | 0.028 | 0.031 | **7.248** | **0.107** | 0.458 | 0.698 | 0.050 | 0.072 | 0.051 | 0.051 |
| D00009.001 | Foxp2 | -0.085 | -0.076 | -0.021 | -0.021 | **10.152** | **0.153** | 0.238 | 0.343 | 0.092 | 0.033 | -0.085 | -0.085 |
| D00011.001 | Irf2 | 0.014 | -0.011 | 0.748 | 0.756 | 0.118 | -0.101 | 4.118 | 4.029 | **0.876** | **10.185** | 0.176 | 0.062 |
| D00619.003 | RFX5 | -0.785 | -0.312 | 3.475 | 1.148 | -1.674 | -1.714 | 2.523 | 2.181 | 1.659 | 1.780 | **5.132** | **-1.959** |
| D00072.001 | Rfx7 | 0.107 | 0.113 | **11.192** | **3.280** | -0.198 | -0.241 | -0.174 | -0.215 | 0.077 | 0.067 | **30.895** | **0.021** |

**Supplementary Figure S3: Non-coding RNA transcription overlapping (A) rs113297984 (*PTPRK – THEMIS*) and (B) rs142852921 (*SKAP2*) for T1D.**

**A: rs113297984**

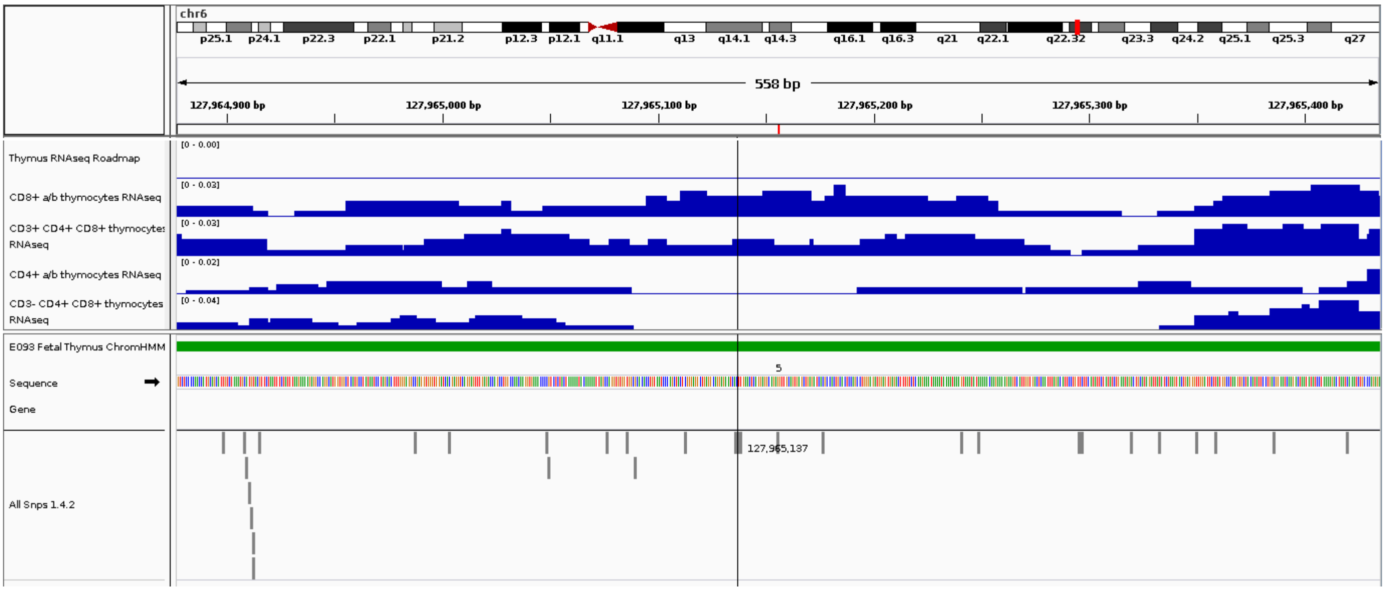

**B: rs142852921 (SKAP2)**

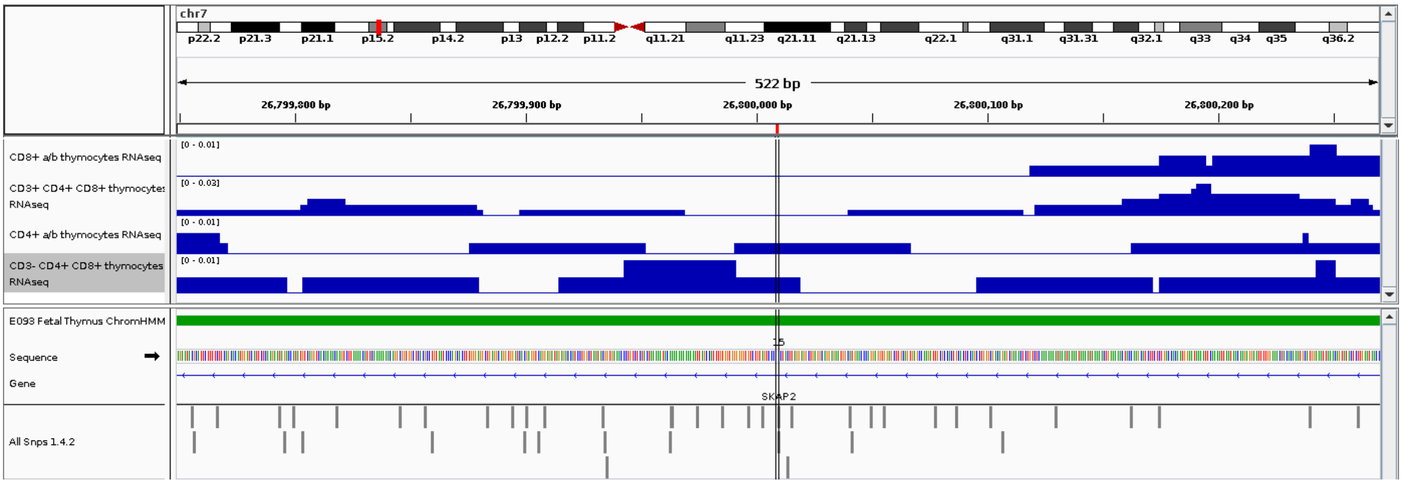
